# NeoPrecis: Enhancing Immunotherapy Response Prediction through Integration of Qualified Immunogenicity and Clonality-Aware Neoantigen Landscapes

**DOI:** 10.1101/2025.07.23.666355

**Authors:** Ko-Han Lee, Timothy J. Sears, Maurizio Zanetti, Hannah Carter

## Abstract

Despite the transformative impact of cancer immunotherapy, the need for improved patient stratification remains critical due to suboptimal response rates. While neoantigens are central to anti-tumor immunity, current metrics like tumor mutation burden are limited by their neglect of immunogenicity and tumor heterogeneity. We present NeoPrecis, a computational framework designed to refine neoantigen characterization across MHC-I and MHC-II pathways and integrate tumor clonality to improve immunotherapy response prediction. NeoPrecis features an interpretable T-cell recognition model that reveals the critical influence of MHC molecules on TCR recognition beyond mere antigen presentation. Benefit HLA alleles identified through model-driven contribution analysis exhibit significant predictive power for patient outcomes in immune checkpoint inhibitor treatment (melanoma: p-value = 0.04; NSCLC: p-value = 0.01). Applying NeoPrecis to immunotherapy-treated tumors, we show the clonality-aware neoantigen landscape improves response prediction in melanoma and heterogeneous NSCLC, achieving 11% and 20% improvement of AUROC compared to TMB respectively. Heterogeneous NSCLCs, more common among never smokers, retain more subclonal neoantigens due to lower immunoediting pressure, where NeoPrecis better captures the varying prevalence of neoantigens. We propose NeoPrecis as a more comprehensive evaluative framework for neoantigen assessment by incorporating both immunogenicity and tumor clonality, offering insights into the link between collective quality of neoantigen landscapes and immunotherapy response.

## INTRODUCTION

Immunotherapy has revolutionized cancer treatment, with personalized cancer vaccines and immune checkpoint inhibitors (ICIs) emerging as two major modalities targeting tumor-specific neoantigens^1,2^. Cancer vaccines work by boosting the immune system through the expansion of T cell precursors specific for these tumor-specific antigens. Since 2017, numerous cancer vaccine trials have been conducted in cancers such as melanoma^3–6^, glioblastoma^7,8^, pancreatic cancer^9^, and others^6,10,11^. However, response rates remain suboptimal. For instance, Rojas *et al.* reported an overall response rate of 50%, yet only 11% of target neoantigens successfully induced a T cell response^9^. These limitations underscore the need for improved strategies to identify and target immunogenic neoantigens more effectively.

Meanwhile, ICIs have demonstrated significant efficacy in certain cancer types, such as melanoma^12,13^ and non-small cell lung cancer (NSCLC)^14,15^, by blocking immune checkpoint pathways and reinvigorating CD8+ T cells recognizing tumor-specific antigens, including neoantigens. Despite their transformative potential, ICIs exhibit variable and often suboptimal response rates^16,17^. For instance, in melanoma, one of the most responsive cancer types, single-agent ICI achieves a response rate of approximately 40%, while combination therapy increases this to around 60%^18,19^. However, some patients experience life-threatening immune-related adverse events after receiving ICIs^20^. Tumor mutation burden (TMB), an FDA-approved biomarker for immunotherapy, acts as a proxy for immunogenic neoantigen abundance^21^. It is known to correlate with response rates across various cancer types^16^. High-TMB cancers like melanoma show higher response rates, while low-TMB cancers such as sarcoma have lower efficacy. Nevertheless, within individual cancer types, TMB’s predictive power is limited^17^. Tumor neoantigen burden (TNB), which relates to the count of MHC-presented neoantigens, offers a more refined metric of the tumor’s immune landscape but fails to account for T-cell receptor (TCR) recognition or the clonal distribution of mutations within tumors^22^.

Neoantigens are novel peptide antigens derived from tumor-specific mutations, playing a critical role in the immune system’s ability to recognize and target tumors. The immunogenicity of neoantigens is highly associated with their abundance and peptide binding affinity for the major histocompatibility complex (MHC) which displays them on the cell surface^23^. However, these factors alone are insufficient for a sensitive and efficient selection of neoantigens^24^. The activation of T cells requires T-cell receptor (TCR) recognition of the peptide-MHC (pMHC) complex. Several indirect metrics have been proposed to evaluate TCR recognition, including agretopicity ratio (the ratio of MHC-binding affinity between mutated and wild-type peptides)^23,25^ whereby a mutant peptide could be more easily recognized by the TCR repertoire if the wild-type counterpart was not involved in thymic selection, and foreignness (measuring the sequence similarity to foreign immunogenic antigens)^26^. Direct approaches include ICERFIRE^27^, DeepNeo^28^ and PRIME^29,30^, which predict immunogenicity based on peptide sequence, and cross-reactivity distance^31^, which measures the distance between wild-type and mutated peptides in terms of TCR recognition. Despite these efforts, the efficacy of TCR recognition metrics remains limited due to the high diversity of TCR repertoires, shaped by genetic recombination and thymic selection^32,33^. Additionally, MHC allele variability across individuals leads to distinct peptide repertoires, further constraining the generalizability of predictive metrics, underscoring the need for improved methods.

Tumor clonality further shapes the immune response by influencing the distribution of neoantigens across tumor subclones. Clonal neoantigens, which are derived from mutations shared by all tumor cells, can generate more effective clinical response than subclonal neoantigens, which are confined to a subset of tumor cells^34,35^. Highly heterogeneous tumors, marked by a greater proportion of subclonal mutations, may evade immune surveillance due to reduced clonal neoantigen exposure^36–38^. Thus, CD8+ T cells predominantly react to clonal mutations in early-stage NSCLC^34^, and NSCLC tumors with a high proportion of subclonal copy-number alterations are at higher risk for recurrence or death than those with a low proportion^39^. Likewise, high tumor heterogeneity correlates with limited ICI response in both mice and humans in melanoma^40^ and with poor survival outcomes in a meta-analysis of the TCGA cohort^38^. These findings suggest that suboptimal clinical outcomes may reflect the inability of the immune cells to cope with tumor heterogeneity and underscore the importance of identifying immunogenic neoantigens, as they are not only important biomarkers of ICI response but also indicators of tumor-immune co-evolution through immune selection.

Historically, neoantigen discovery has predominantly focused on the MHC-I pathway^41^, which activates CD8+ T cells for direct tumor cell killing. This focus reflects the established role of CD8+ T cells as primary cytotoxic effectors in anti-tumor immunity. However, recent studies have expanded our understanding of CD4+ T cells, highlighting their critical roles in both direct tumor killing and CD8+ T cell priming. Bawden *et al.* revealed direct cytotoxic effects of CD4+ T cells against melanoma^42^, while Espinosa-Carrasco *et al.* emphasized their role in sustaining CD8+ T cell responses through interactions with intra-tumor dendritic cells (DCs)^43^. Further supporting this, Pyke *et al.*^44^ and Alspach *et al.*^45^ demonstrated the contribution of MHC-II neoantigens to anti-tumor immunity. Moreover, Sears *et al.*^46^ found that patients with MHC-II-reliant neoantigen profiles, featuring a higher proportion of MHC-II neoantigens, experience significantly longer-lasting clinical benefits. These findings reinforce the critical role of MHC-II neoantigens and highlight the necessity of identifying both MHC-I and MHC-II neoantigens to fully capture the cooperative interplay between CD8+ and CD4+ T cells in tumor immunity.

To encompass these variables, we developed NeoPrecis, a computational framework that predicts neoantigen immunogenicity and constructs a clonality-aware neoantigen landscape model to more accurately evaluate patients’ potential immune responses (**Fig. 1**). NeoPrecis integrates three critical dimensions—neoantigen abundance, MHC presentation, and TCR recognition—into a unified evaluative framework. This framework features an interpretable immunogenicity model that accounts for MHC allele variability, enhancing the accuracy of TCR recognition predictions. By incorporating tumor clonality into the neoantigen landscape model, NeoPrecis also addresses limitations of current metrics like TMB and TNB, providing a more precise representation of the tumor’s immune landscape. Additionally, NeoPrecis incorporates both MHC-I and MHC-II pathways to provide a detailed representation of tumor-immune interactions. This comprehensive approach improves the identification of immunogenic neoantigens and enables better patient stratification for immunotherapy.

**Fig. 1.**
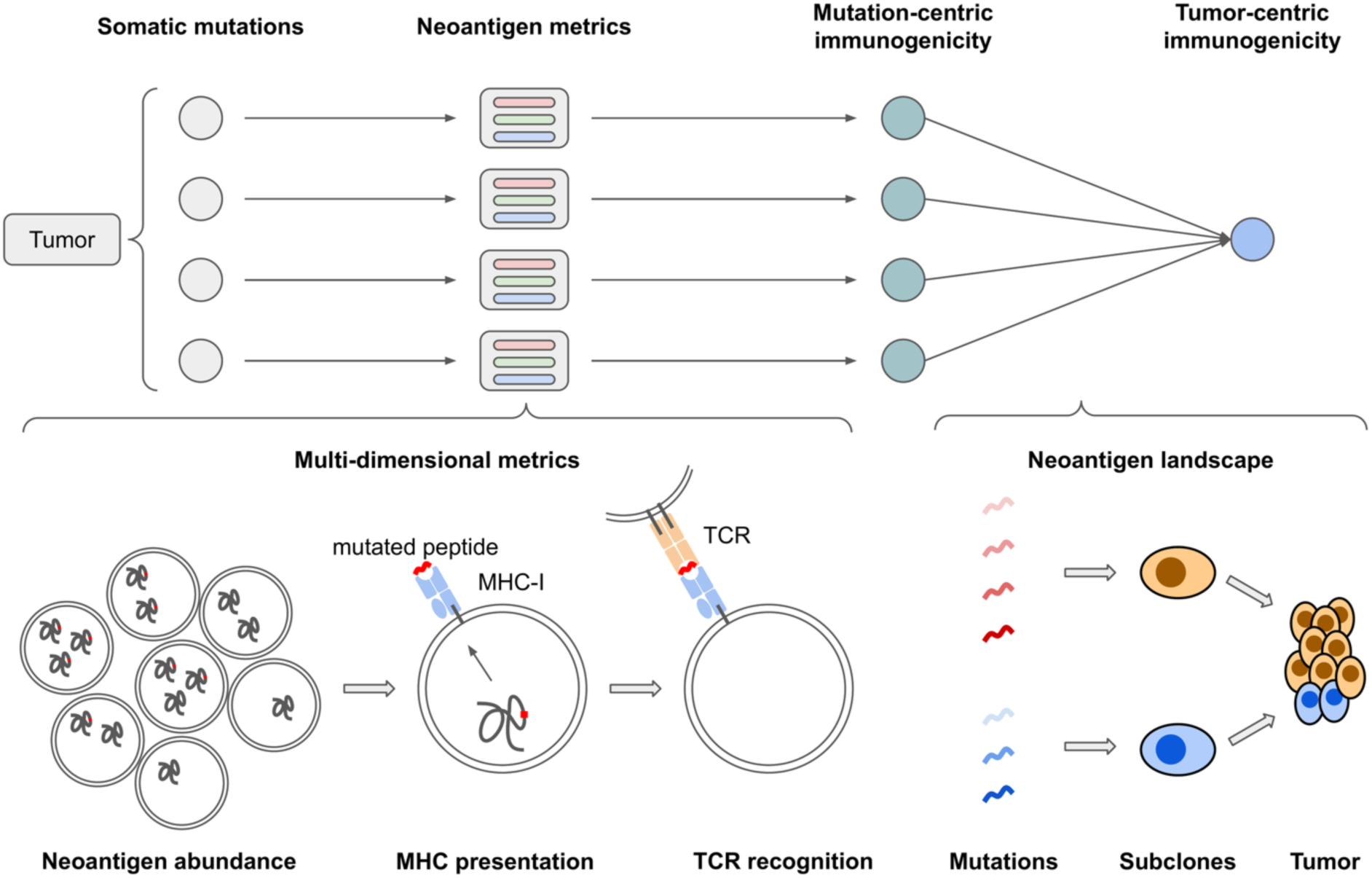
Overview of NeoPrecis for tumor neoantigen assessment. The workflow begins with the computation of neoantigen-related multi-dimensional metrics, including neoantigen abundance, MHC presentation, and TCR recognition, for each protein-altering somatic mutation. These metrics are subsequently integrated to derive mutation-centric immunogenicity scores. Finally, a tumor-centric immunogenicity score is generated using a framework designed to map the neoantigen landscape considering neoantigen immunogenicity and tumor clonality.

## RESULTS

### Characterizing T-cell recognition in immunogenicity prediction

Measures for assessing neoantigen abundance and binding to MHC molecules are well established. In contrast, predicting T-cell recognition is challenging due to the high variability of TCR repertoires and the still limited availability of TCR sequencing data. To incorporate a model of T cell recognition into our *in silico* neoantigen prioritization pipeline, we considered two alternative approaches for assessing TCR recognition without TCR sequencing: peptide-based methods and the cross-reactivity distance (CRD) approach. Cross-reactivity, a well-documented feature of TCR-peptide binding, describes the ability of a TCR to bind multiple peptides with similar sequences^47^. Peptide-based methods, such as PRIME^29,30^ and DeepNeo^28^, primarily model the immunogenicity of target peptides. In contrast, CRD, first introduced by Łuksza *et al.*^31^, measures the differences between wild-type (WT) and mutated (MT) peptides, aligning more closely with immune mechanisms, as thymic selection acts to constrain TCR diversity to distinguish self from non-self peptides^33^. However, existing models overlook the role of MHC-binding motifs, which are critical in shaping TCR-pMHC interactions. To address this limitation, we developed a new immunogenicity model for NeoPrecis, NeoPrecis-Immuno, which incorporates MHC-binding motifs into a refined CRD-based framework, enabling a more nuanced characterization of neoantigens.

NeoPrecis-Immuno embeds MHC-binding core regions (9-mer) using trained amino acid embeddings and MHC-binding motif enrichment, projecting each residue into a latent space specific to MHC alleles. Residue-level embeddings are then aggregated using position-weighted pooling to generate peptide embeddings for WT and MT peptides, highlighting the varying importance of core positions in TCR-pMHC interactions. To capture differences between WT and MT peptides, NeoPrecis-Immuno employs a geometric representation that includes origin, direction, and distance. The distance component predominantly reflects CRD and is further refined with peptide sequence information (origin and direction). Finally, NeoPrecis-Immuno integrates CRD with MHC-binding score as a covariate, allowing the model to further learn immunogenicity beyond what is captured by MHC binding alone, to output a probability that represents neoantigen immunogenicity (**Fig. 2A**).

**Fig. 2.**
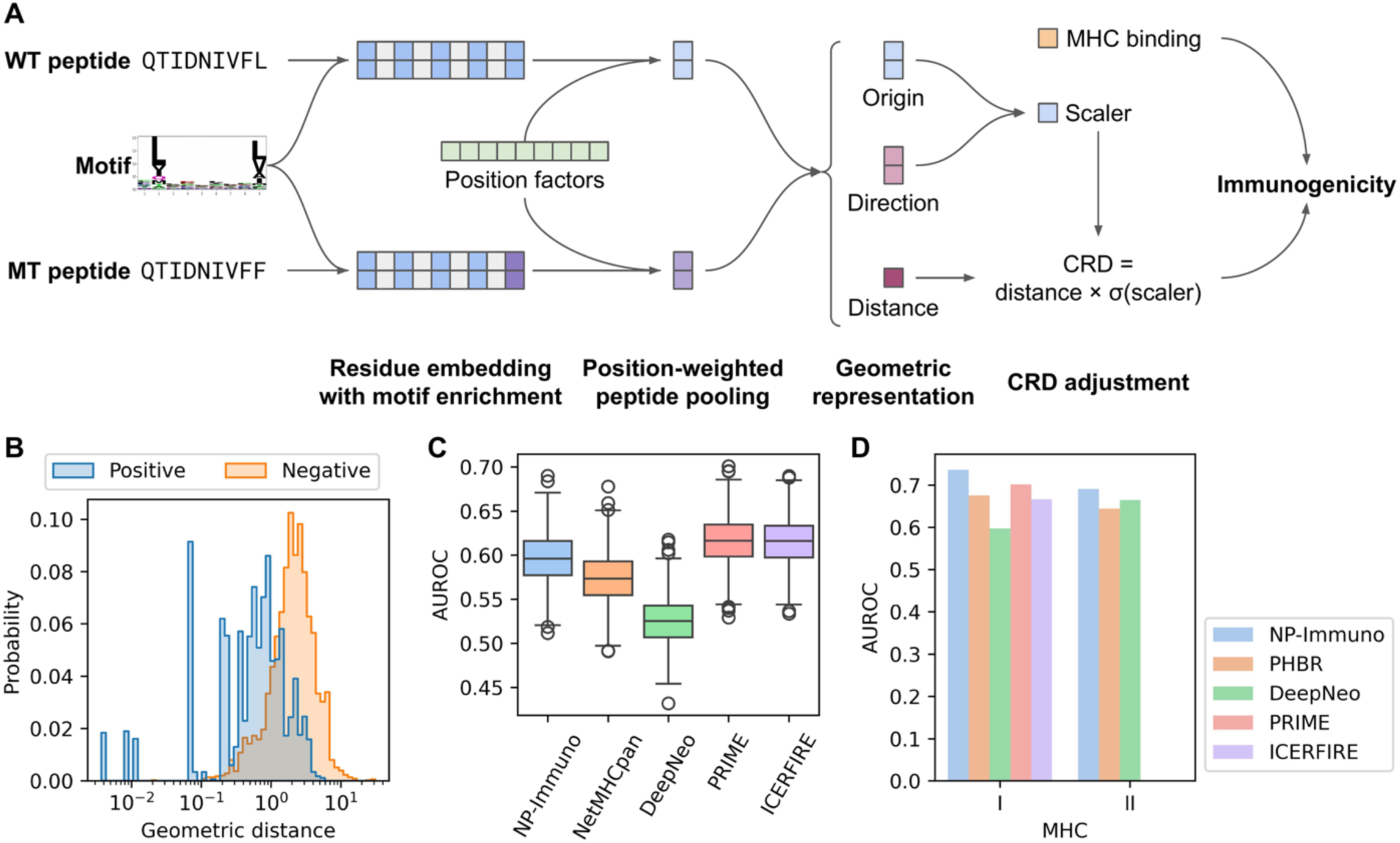
Development and validation of the T-cell recognition model, NeoPrecis (NP)-Immuno. **(A)** Schematic of NeoPrecis-Immuno architecture. Peptide residues are embedded with motif enrichment features, and their contributions are weighted by trainable position factors to generate a peptide embedding. A geometric representation captures the differences between wild-type (WT) and mutated (MT) peptides, from which the cross-reactivity distance is computed. Combined with the MHC binding prediction as a covariate, the model outputs an immunogenicity score highlighting T-cell recognition potential. **(B)** Distribution of geometric distances in the TCR-binding triplet dataset. The positive group consists of positive pairs (seed and positive peptides) that mimic non-immunogenic substitutions, while the negative group consists of negative pairs (seed and negative peptides) that mimic immunogenic substitutions. **(C)** Area under the receiver operating characteristic curve (AUROC) for each predictor on MHC-I samples (n = 438; #positives = 228) from the internal testing set (CEDAR dataset). AUROC distributions were calculated using bootstrapping (k = 1,000). In the boxplots, the center line represents the median, boxes indicate the interquartile range (IQR; 25th–75th percentile), and whiskers extend to the most extreme data points within 1.5× the IQR. Outliers are depicted as individual circles. **(D)** AUROC comparisons of each predictor on MHC-I (n = 1,088; #positives = 36) and MHC-II (n=1,036; #positives = 30) samples from the external testing set (NCI dataset). PHBR is an allele-aggregating score for MHC-binding prediction.

We implemented a two-stage training process using TCR-pMHC binding data from IEDB^48^ and VDJdb^49^ (**Table S1**) and T-cell assay data from CEDAR^50^ (**Table S2**). In the first stage, we constructed a triplet dataset from binding data, where each triplet consists of a seed peptide (mimicking a wild-type peptide), a positive peptide (mimicking a non-immunogenic mutated peptide), and a negative peptide (mimicking an immunogenic mutated peptide). The model was trained to minimize the distance between positive pairs (seed and positive) and maximize the distance between negative pairs (seed and negative), simulating the distinction between non-immunogenic and immunogenic neoantigens (**Fig. S1A**, **Methods**). Notably, the geometric distance calculated by NeoPrecis-Immuno outperformed the BLOSUM62 distance in differentiating positive and negative peptide pairs (**Fig. 2B**, **Fig. S2A**). In the second stage, we fine-tuned the model using T-cell assay data with T-cell activation labels, enabling it to learn the immunogenicity of target peptides (**Methods**). The resulting immunogenicity model was subsequently validated on both an internal and an external testing set to benchmark its performance.

We compared NeoPrecis-Immuno to an MHC-binding predictor (NetMHCpan^51,52^) and immunogenicity models (PRIME^29,30^, ICERFIRE^27^, and DeepNeo^28^) (**Methods**). The internal testing set, isolated from the CEDAR dataset^50^ before training, is epitope-based with labels for each pMHC, while the external testing set, derived from an independent cancer cohort (NCI dataset, Parkhurst *et al.*^53^), is mutation-based with T-cell assay data for each mutation (**Table S3**). For mutation-based predictions, aggregation across MHC alleles was required (**Methods**). As there weren’t many MHC-II datapoints, internal validation was performed exclusively for MHC-I. To ensure fairness, we identified pMHCs not present in the training datasets of any predictors (n = 1,147, **Fig. S2B**) and excluded pMHCs with unavailable MHC alleles in any predictor. Among the remaining 438 pMHCs (#positives = 228), NeoPrecis-Immuno outperformed NetMHCpan and DeepNeo but performed slightly worse than PRIME and ICERFIRE (**Fig. 2C**, **Fig. S2C**). For external validation, NeoPrecis-Immuno achieved the best performance for both MHC-I (n = 1,088; #positives = 36) and MHC-II (n = 1,036; #positives = 30) predictions (**Fig. 2D**, **Fig. S2D**). Although mutations with no available alleles were excluded, for some mutations, not all alleles were supported (< 6 alleles for MHC-I; < 10 alleles for MHC-II) by ICERFIRE and DeepNeo (**Fig. S2E**), potentially contributing to their lower performance. Overall, these benchmarks demonstrate that NeoPrecis-Immuno is competitive with other immunogenicity predictors.

### Interpretable MHC-binding motif contributions to T-cell recognition

To evaluate the contribution of each component in NeoPrecis-Immuno, we extracted scores from the model for substitution distance (SubDist), substitution distance with position weighting (SubPosDist), geometric distance (GeoDist; SubPosDist with motif enrichment), and cross-reactivity distance (CRD; GeoDist with sigmoid scaling) (**Table S4**, **Methods**). BLOSUM62 distance (BLOSUMDist) was included as a baseline for comparison. These distances quantify the difference between WT and MT peptides. All NeoPrecis-Immuno components outperformed the baseline (BLOSUMDist) in both MHC-I and MHC-II contexts (**Fig. 3A**). For MHC-I, performance steadily improved as additional components were added to the model. However, in MHC-II, differences between components were less pronounced, likely due to the smaller sample size. Sigmoid scaling contributed minimally to CRD, as GeoDist and CRD scores were highly correlated (**Fig. S3A**).

**Fig. 3.**
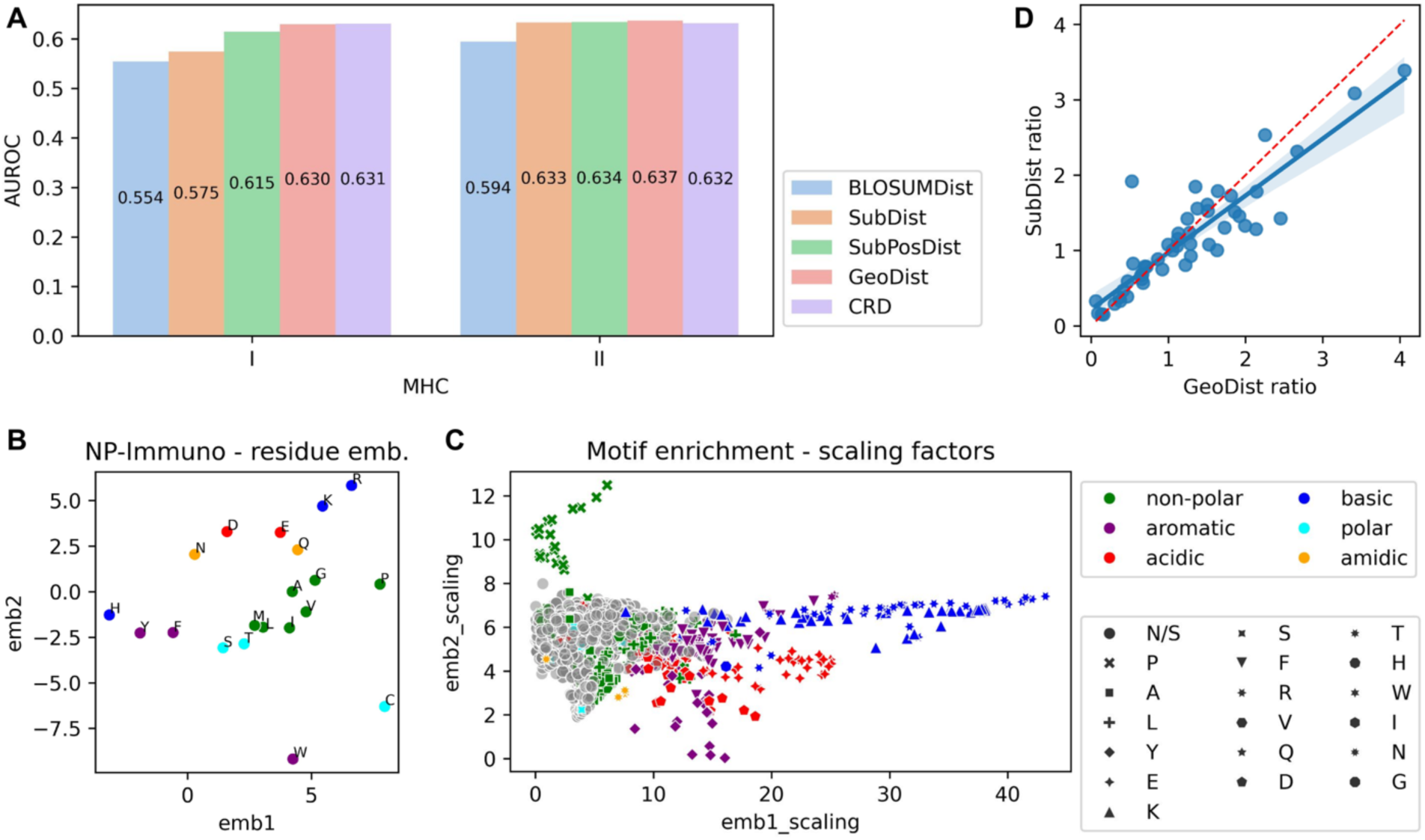
Interpretation of NeoPrecis-Immuno. **(A)** AUROC comparisons of incrementally constructed NeoPrecis-Immuno components for MHC-I and MHC-II predictions. Each component builds upon the previous one, except for BLOSUMDist, which serves as a baseline comparison. The components include: BLOSUMDist (BLOSUM62 distance, baseline), SubDist (substitution distance of residue embeddings), SubPosDist (SubDist with position weighting), GeoDist (SubPosDist with MHC-binding motif enrichment), and CRD (GeoDist with sigmoid scaling). **(B)** Distribution of residue embeddings in NeoPrecis-Immuno. Residues are colored by amino acid properties to illustrate clustering based on biochemical characteristics. **(C)** Scaling factors for motif enrichment across MHC allele positions. These factors indicate how motif enrichment adjusts residue embeddings (B) along specific axes. Each dot is annotated with the most frequently observed amino acid at the binding motif for that allele position, represented by both color and text. **(D)** Positive-to-negative ratio distributions for SubDist and GeoDist across MHC-I allele-position pairs in the NCI dataset. Positives represent immunogenic substitutions while negatives represent non-immunogenic substitutions. A higher ratio indicates better differentiation. The red line represents x = y, while the blue line shows the fitted regression.

Plotting amino acid embeddings on a 2D projection, NeoPrecis-Immuno residue embeddings grouped amino acids based on their chemical properties, with similar residues clustering together (**Fig. 3B**). Notably, NeoPrecis-Immuno embeddings differed from BLOSUM62 embeddings; for instance, histidine (H) was positioned closer to tyrosine (Y) and phenylalanine (F), likely due to their shared aromatic rings, while tryptophan (W) was positioned further away, possibly reflecting its larger size (**Fig. S3B**). Thus, NeoPrecis-Immuno embeddings obtained through training on TCR-pMHC interaction data differ from BLOSUM62 embeddings, derived from evolutionary conservation more broadly.

Position factors revealed that the third and fifth residue position were most influential for MHC-I and MHC-II, respectively (**Fig. S3C**). Interestingly, these are not typical anchor positions for MHC binding, suggesting their significance lies in TCR-pMHC interactions. This hypothesis is supported by the TCR-binding triplet dataset, where amino acid substitutions with significant property changes (BLOSUM62 substitution score ≤ 0) at these positions were rare (**Fig. S1B**).

The motif enrichment step of NeoPrecis-Immuno played a critical role in refining residue embeddings. Using affine transformations (rotation, reflection, and scaling) for interpretation, we analyzed how motif-enriched residue embeddings align with MHC-binding motifs (**Methods**). Among these transformations, scaling was the primary contributor to differences between wild-type and mutated amino acids, with scaling factors being specific to each position and allele. This specificity likely reflects the variation in MHC-binding motifs across different positions. Positions dominated by the same amino acid exhibited consistent scaling behavior. For example, positions dominated by proline tended to scale the y-axis, while those dominated by arginine scaled the x-axis (**Fig. 3C**). These refinements adjusted the relative distances between amino acids in the embedding space, improving the ability of NeoPrecis-Immuno to differentiate wild-type and mutated residues for immunogenic neoantigen prediction.

To quantify the impact of motif enrichment, we analyzed the positive-to-negative distance ratio for different metrics, where positive distance indicates the distance between a wild-type peptide and its immunogenic mutated counterpart, while negative distance indicates the distance between a wild-type peptide and its non-immunogenic mutated counterpart. Neoantigens were grouped by allele and position to minimize bias, and the mean distances for positive and negative pairs were used to compute the ratio. GeoDist demonstrated a larger positive-to-negative ratio compared to SubDist (**Fig. 3D**), revealing that motif enrichment enhanced the model’s ability to capture meaningful distance relationships between amino acids and better distinguish immunogenic from non-immunogenic samples.

We further computed an allele benefit score by averaging scaling factors across positions by MHC molecule (**Table S5**, **Methods**). A higher benefit score indicates greater overall scaling of amino acid distance, reflecting increased immunogenicity and potential benefits for tumor immunity. To analyze trends, we categorized alleles based on the most dominant amino acid at the anchor position of their binding peptides, which contributes most to the allele benefit score. MHC-I alleles associated with glutamic acid (E), lysine (K), or arginine (R) exhibited the highest benefit scores (**Fig. S3D**), aligning with previously reported benefit HLA supertypes, including HLA-B27, which predominantly binds peptides with arginine at the anchor position, and HLA-B44, which favors peptides with glutamic acid^54,55^.

### Multi-dimensional metrics for neoantigen immunogenicity

Beyond TCR recognition, neoantigen abundance and MHC presentation are two major factors contributing to immunogenicity. We defined eight key features across the three dimensions and evaluated these dimensions with the NCI dataset^53^ (**Methods**). For neoantigen abundance, DNA allele frequency (AF) serves as a rough approximation of mutation clonality^56^, while RNA AF and RNA expression levels quantify the expression of mutated peptides. To capture MHC presentation, patient harmonic-mean best rank (PHBR) measures binding affinity across an individual’s MHC alleles^57^, and robustness indicates the number of binding MHC alleles. For TCR recognition, in addition to the NeoPrecis-Immuno score, we included agretopicity ratio, proposed by Duan *et al.*^25^, and foreignness, proposed by Łuksza *et al.*^26^.

While PHBR, which aggregates netMHCpan predictions across MHC alleles using a harmonic mean, has shown robust mutation-centric performance for predicting cell surface presentation^57^, no standard method exists for aggregating TCR recognition metrics across alleles. To address this, we tested several aggregation strategies (**Fig. S4**, **Methods**); The masked maximum approach, which prioritizes the peptide with the highest recognition score across alleles that meet a minimum binding threshold necessary for presentation, either outperformed or matched the best binding approach across all three recognition metrics when evaluated on T-cell activity (**Fig. 4A**). Unlike the best binding approach, which focuses solely on the MHC allele with the highest binding prediction, the masked maximum approach considers all potential binding alleles. These results suggest that TCR recognition is best captured by considering all alleles capable of effective presentation, rather than focusing solely on the strongest binding peptide.

**Fig. 4.**
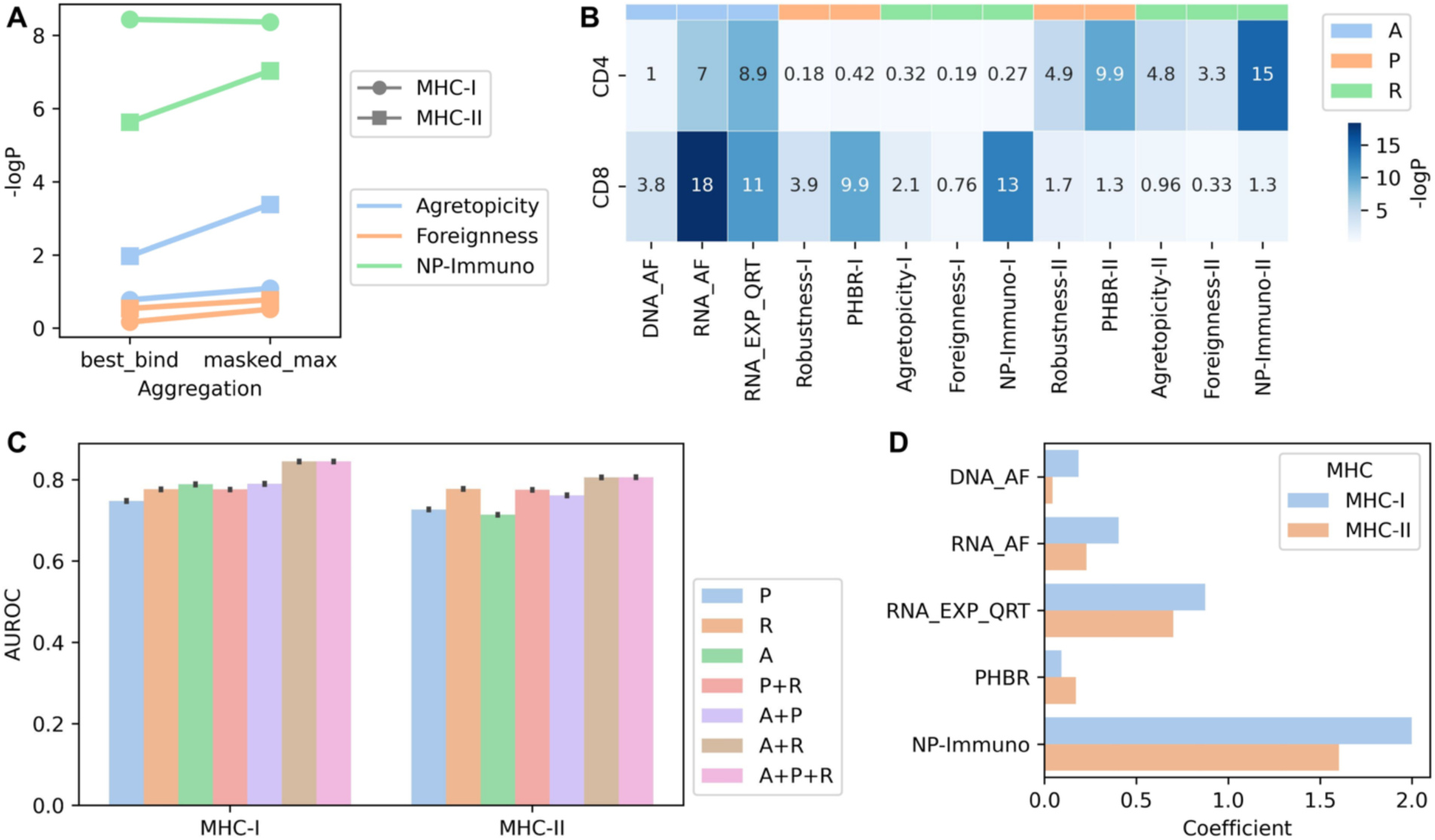
Integration of multi-dimensional metrics for immunogenicity prediction. **(A)** Comparison of allele aggregation methods. Results of the Mann-Whitney U test assess the association between TCR recognition metrics and T cell activities. “Best_bind” indicates the best-binding allele approach, while “masked_max” refers to the masked maximum approach. **(B)** Association test for individual metrics in the NCI cohort. Mann-Whitney U test results evaluate the relationship between each metric and T cell activities, where “NP” denotes “NeoPrecis”, “I” represents MHC-I, “II” represents MHC-II, “A” denotes abundance metrics, “P” denotes presentation metrics, and “R” denotes recognition metrics. “RNA_EXP_QRT” denotes the quartile of RNA expression level (TPM). Agretopicity denotes the agretopicity ratio. **(C)** AUROC values for cross-validation on the NCI cohort using different combinations of metrics. **(D)** Feature importance in the logistic regression model integrating multi-dimensional metrics.

We next assessed the association between mutation-centric features and T-cell activity. Consistent with established immune mechanisms, MHC-I metrics were primarily associated with CD8+ T-cell activity, while MHC-II metrics correlated with CD4+ T-cell activity (**Fig. 4B**). Among the metrics, RNA AF, RNA expression, PHBR, and NeoPrecis-Immuno emerged as the strongest predictors of immunogenicity. DNA AF showed particular importance for MHC-I, while agretopicity ratio and foreignness were more influential for MHC-II predictions.

To refine predictions, we integrated the mutation-centric features into a logistic regression model. After performing feature selection, DNA AF, RNA AF, RNA expression, PHBR, and NeoPrecis-Immuno were included in the final integrated multi-dimensional model, NeoPrecis-Integrated (**Fig. S5A**, **Methods**). Cross-validation on the NCI dataset demonstrated that the three-dimensional model outperformed single- and two-dimensional models (**Fig. 4C**, **Fig. S5B**). Feature importance analysis revealed that NeoPrecis-Immuno was the most significant metric in the integrated model of both MHC-I and MHC-II (**Fig. 4D**). Thus, integrating multi-dimensional mutation-centric features improved neoantigen immunogenicity prediction, with NeoPrecis-Immuno emerging as the most influential feature.

### Advancing immunotherapy response analysis with NeoPrecis

TMB and TNB are designed to quantify the number of potentially immunogenic neoantigens but often underperform due to their limited consideration of immunogenic factors. To enhance neoantigen identification, we applied NeoPrecis to a meta ICI cohort comprising five melanoma^58–62^ and three non-small cell lung cancer (NSCLC)^35,63,64^ cohorts (**Table S6**, **Methods**). This dataset included 695 samples with whole-exome sequencing data. We filtered for patients with pre-treatment biopsies and no prior ICI treatment to avoid pre-existing immune effects (**Fig. 5A**). One sample that failed clonal analysis and nine samples lacking RECIST labels were excluded, resulting in 525 patients (277 melanoma and 248 NSCLC). For subsequent analyses, ICI response was defined as complete response (CR) and partial response (PR) while non-response was defined as stable disease (SD) and progressive disease (PD). Overall survival (OS) was assessed for melanoma, while progression-free survival (PFS) was evaluated for NSCLC due to data availability (**Table S7**).

**Fig. 5.**
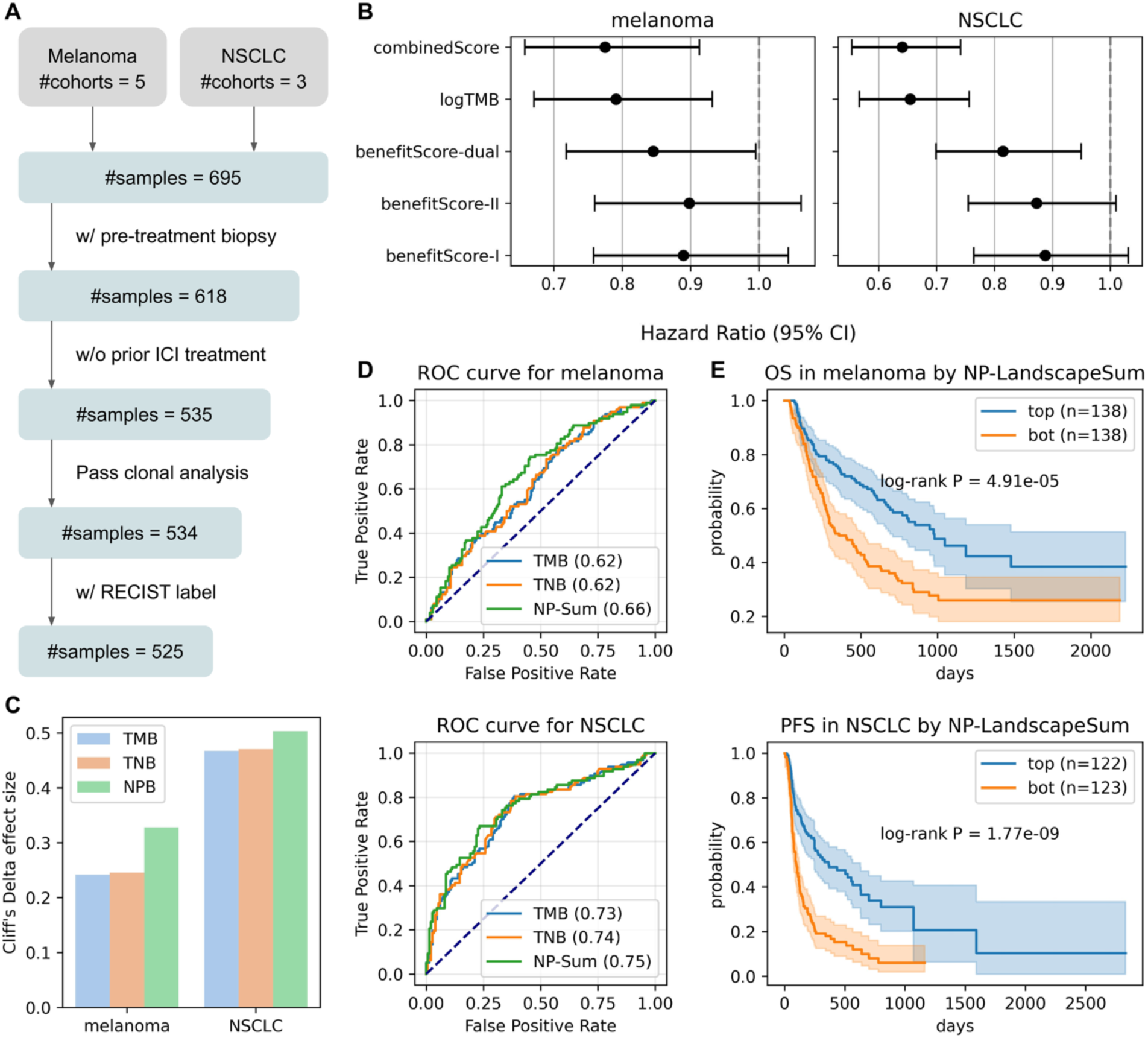
Prediction of ICI response using the neoantigen landscape model. **(A)** Criteria for patient inclusion in the analysis. **(B)** Hazard ratio of metrics related to allele benefit scores derived from the Cox proportional hazards model with sex and age as confounders. Overall survival (OS) is used for melanoma, while progression-free survival (PFS) is used for NSCLC. “Dual” is the metric considering both MHC-I and MHC-II. “CombinedScore” is the product of “logTMB” and “BenefitScore-dual”. **(C)** Cliff’s Delta effect sizes comparing the impact of different mutation burdens on ICI response, including tumor mutation burden (TMB), tumor neoantigen burden (TNB), and NeoPrecis-Immuno burden (NPB), for melanoma (n = 277; #positives = 98) and NSCLC (n = 248; #positives = 97). **(D)** AUROC comparison of ICI response prediction using TMB, TNB, and NeoPrecis-LandscapeSum (NP-Sum) in melanoma and NSCLC. **(E)** Kaplan-Meier survival curves stratified by NP-LandscapeSum, with log-rank P-values for melanoma (OS) and NSCLC (PFS). “Top” refers to the top half of patients with higher NP-LandscapeSum scores, while “bot” refers to the bottom half of patients with lower NP-LandscapeSum scores. NP denotes NeoPrecis.

First, we assessed the allele-level contribution to ICI treatment outcomes using the allele benefit score, derived from the NeoPrecis-Immuno model, which quantifies how MHC alleles refine neoantigen immunogenicity. A patient-centric allele benefit score (BenefitScore) was calculated as the geometric mean across all alleles (**Methods**). We then applied a Cox proportional hazards model, incorporating age and sex as confounders, to estimate its prognostic value. BenefitScore-dual, which integrates both MHC-I and MHC-II, was statically significant in predicting patient outcomes for both melanoma (p-value = 0.04) and NSCLC (p-value = 0.01), and outperformed BenefitScore-I (MHC-I only) and BenefitScore-II (MHC-II only) (**Fig. 5B**). We further compared its performance against TMB as a baseline. While BenefitScore alone did not surpass TMB, their combination (CombinedScore) improved the overall predictive power (**Methods**). These findings support that different MHC alleles modulate neoantigen immunogenicity in distinct ways, ultimately influencing ICI treatment outcomes.

Since allele-level assessment doesn’t directly account for mutations, we next evaluated the mutation-level contribution using NeoPrecis-Immuno burden (NPB), defined as the number of mutations passing an NeoPrecis-Immuno threshold of 0.4 (approximately the median) for both MHC-I and MHC-II (**Methods**). NPB demonstrated superior performance in stratifying immune responses compared to TMB and TNB, as evidenced by both Cliff’s Delta effect size (**Fig. 5C**) and a non-parametric statistical test (**Fig. S6A**), suggesting that NeoPrecis-Immuno improves neoantigen immunogenicity predictions by accounting for TCR recognition.

To conduct a more comprehensive mutation-level evaluation, we calculated a tumor-centric immunogenicity score, NeoPrecis-LandscapeSum, by summing NeoPrecis-Immuno predictions across all mutations (**Methods**). For each mutation, the NeoPrecis-LandscapeSum MHC-dual score was derived by multiplying the MHC-I and MHC-II NeoPrecis-Immuno predictions. This integrated approach outperformed NeoPrecis-LandscapeSum MHC-I-only or MHC-II-only scores (**Fig. S6B**, **Methods**) and demonstrated superior performance over TMB and TNB in both melanoma (AUROC = 0.66) and NSCLC (AUROC = 0.75) (**Fig. 5D**). Given the better performance of MHC-dual, we refer to the tumor-centric immunogenicity score as MHC-dual in subsequent analyses. In survival analyses (**Methods**), patients in the top half of the tumor-centric score showed significantly better outcomes than those in the bottom half (**Fig. 5E**). NeoPrecis-LandscapeSum outperformed TMB in melanoma and showed comparable performance in NSCLC (**Fig. S6C**).

Lastly, we evaluated the efficacy of the multi-dimensional integrated immunogenicity prediction model (NeoPrecis-Integrated) in patients with RNA-seq data, identifying 187 samples (137 melanoma and 50 NSCLC). The NeoPrecis-Integrated model, trained on the NCI cohort using five selected features (**Methods**), demonstrated improved performance over NeoPrecis-Immuno in melanoma but not in NSCLC (**Fig. S6D**). Here, the lower purity of NSCLC tumors (**Fig. S6E**) may have impacted the inference of neoantigen abundance metrics, such as DNA AF and RNA AF, implicating the importance of data quality for robust predictions.

### Clonality-aware neoantigen landscapes in tumor heterogeneity

Mutation burden or summation metrics cannot account for the distribution of mutations within tumors. To address this limitation, we sought to characterize the tumor-specific neoantigen landscape by incorporating both immunogenicity and tumor subclonal architecture. First, we performed clonal analysis using PyClone^65^, grouping mutations into clusters annotated with their prevalence in the tumor and mutation number (**Methods**). We developed two approaches to integrate tumor clonality: NeoPrecis-LandscapeCCF, which calculates a weighted sum of NeoPrecis-Immuno scores using cancer cell fraction (CCF), and NeoPrecis-LandscapeClone, which aggregates NeoPrecis-Immuno scores by cluster architecture (**Methods**). For the latter, the NeoPrecis-LandscapeClone scores of mutations within a cluster were summed to determine cluster immunogenicity, which was then averaged across clusters weighted by prevalence to derive tumor-centric immunogenicity (**Fig. 6A**). We benchmarked these against existing approaches that also incorporate mutation clonality within tumors, including the Cauchy-Schwarz index of neoantigens (CSiN)^66^ and the immunoediting-based optimized tumor neoantigen load (ioTNL)^67^. CSiN profiles the neoantigen landscape by weighting the neoantigen load of each mutation with its variant allele frequency, while ioTNL aggregates weighted neoantigen loads across subclones potentially subject to immune elimination (**Methods**). Overall, NeoPrecis outperformed other approaches by more effectively qualifying the immunogenicity of neoantigens within subclones, rather than simply quantifying their number. In melanoma, our clonality-aware methods generally showed better performance with NeoPrecis-LandscapeClone achieving the best AUROC of 0.69. However, in NSCLC, mutation burden-based methods yielded better results, with NeoPrecis-LandscapeSum demonstrating the top AUROC of 0.75 (**Fig. S7A**).

**Fig. 6.**
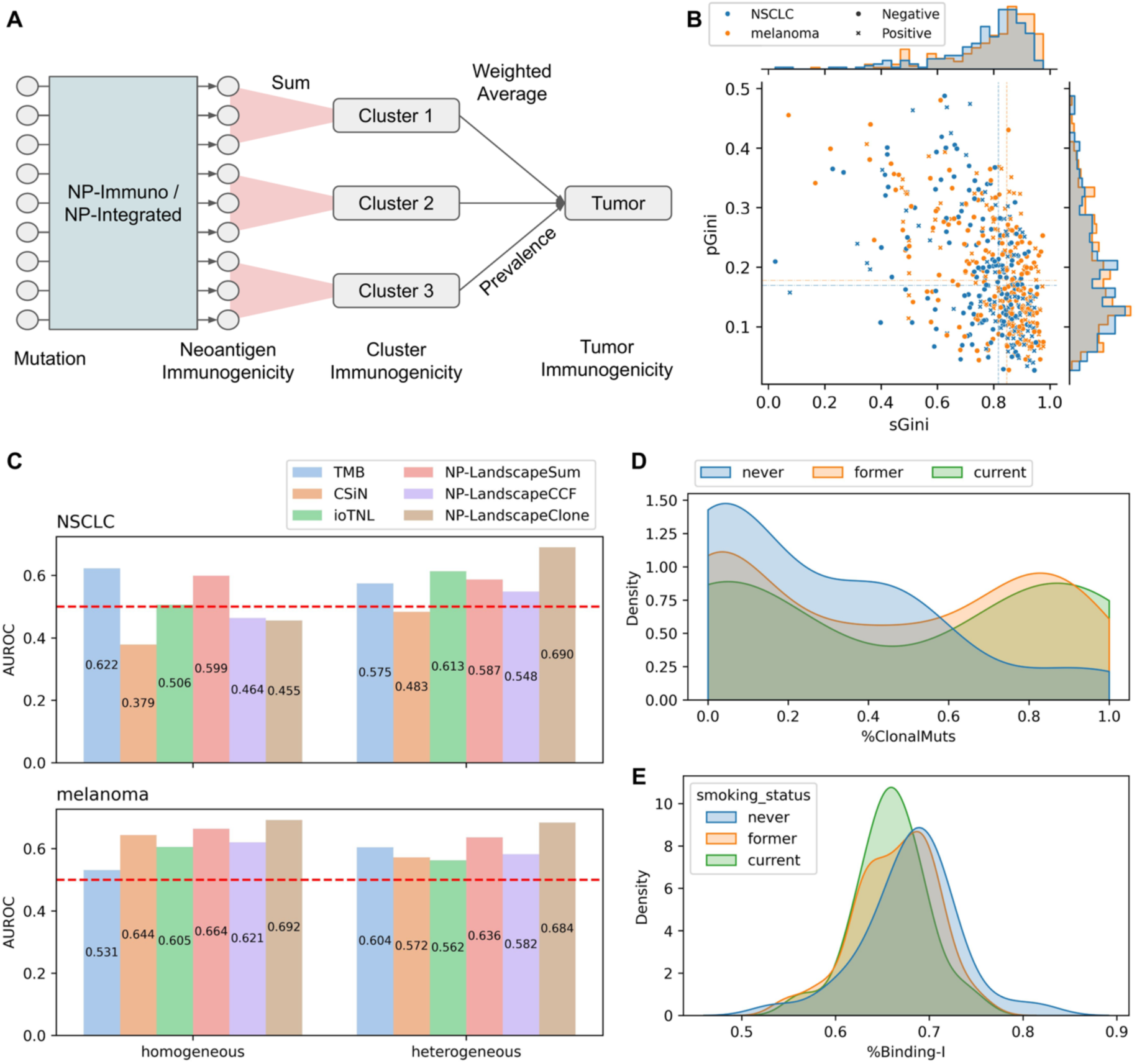
Enhancing ICI prediction by incorporating tumor subclonal architecture. **(A)** Computation of NeoPrecis-LandscapeClone. Immunogenicity predictions from NeoPrecis-Immuno or NeoPrecis-Integrated are summed within each mutation cluster. These cluster sums are then combined using a weighted average, where the prevalence of each cluster serves as the weight. This approach generates a tumor-centric immunogenicity score by integrating multiple mutation-centric predictions. **(B)** Distribution of mutation number Gini index (sGini) and prevalence Gini index (pGini). ICI response (positive or negative) is annotated with the mark shape. **(C)** AUROC comparison of TMB, CSiN, ioTNL, NP-LandscapeSum, NP-LandscapeCCF and NP-LandscapeClone across two patient groups, heterogeneous (low sGini–low pGini) and homogeneous (high sGini–high pGini), in melanoma and NSCLC. In NSCLC, tumor heterogeneity is associated with smoking status, where smokers are more likely to have homogeneous tumors, while never smokers tend to have heterogeneous tumors (**Fig. S8A**). NP denotes NeoPrecis. **(D)** Density plot estimated using kernel density estimation (KDE) showing the distribution of %ClonalMuts, the ratio of clonal mutations (CCF ≥ 0.85) to total mutations, across different smoking statuses. Values are clipped between 0 and 1. **(E)** Density plot estimated using KDE showing the distribution of %Binding-I, the ratio of MHC-I binding mutations (PHBR ≤ 2) to total mutations, across different smoking statuses. Values are clipped between 0 and 1.

To investigate the discrepancy between melanoma and NSCLC, we evaluated tumor heterogeneity for each sample. Tumor heterogeneity was quantified using the Gini index, which measures inequality, across both prevalence and size (mutation number) dimensions to better capture the distribution of mutation clusters (**Methods**). A high Gini index indicates dominance of a few clusters and greater homogeneity. Conversely, a low Gini index reflects a more heterogeneous tumor. Surprisingly, the distributions of prevalence Gini index (pGini) and size Gini index (sGini) were similar between melanoma and NSCLC (**Fig. 6B**). Tumors were then classified based on their heterogeneity: the first quadrant (high pGini and high sGini) represented more homogeneous tumors, while the third quadrant (low pGini and low sGini) represented more heterogeneous tumors. In NSCLC, patients with heterogeneous tumors exhibited worse outcomes compared to those with homogeneous tumors, although the difference was not statistically significant (**Fig. S7B**). In contrast, no such trend was observed in melanoma. We further analyzed tumor heterogeneity in relation to smoking status in NSCLC. Never smokers had a higher proportion of heterogeneous tumors compared to smokers (**Fig. S8A**). Additionally, they exhibited lower TMB (**Fig. S8B**) and are known to have poorer responses to immunotherapy^68^, aligning with the worse prognosis observed in the heterogenous group.

We evaluated the performance of the clonality-aware neoantigen landscape across the heterogeneity-defined groups (**Methods, Fig. 6C, Fig. S7C**). NeoPrecis-LandscapeSum, which does not account for clonality, performed consistently across both groups and tumor types, indicating no subset bias between the groups. In melanoma, NeoPrecis-LandscapeClone consistently outperformed the other methods, regardless of tumor heterogeneity. In NSCLC, however, NeoPrecis-LandscapeClone showed superior performance in the heterogeneous group but underperformed in the homogeneous group. This pattern aligns with the weaker performance of NeoPrecis-LandscapeSum in never smokers (**Fig. S8C**), who tend to have more heterogeneous tumors. These findings suggest that as predicted the clonality-aware neoantigen landscape is particularly effective at capturing immunogenic neoantigens in heterogeneous NSCLC tumors.

The clonality-aware neoantigen landscape primarily reflects the impact of clonal neoantigens; its poor performance in homogeneous NSCLC tumors suggests that clonal neoantigens may play a less significant role in these cases. One possible explanation is that these tumors have undergone stronger immunoediting, leading to the selective elimination of highly immunogenic clonal neoantigens. To investigate this, we examined the proportion of clonal mutations (CCF ≥ 0.85)^35^ and the ratio of MHC-I binding mutations (PHBR ≤ 2) across different smoking statuses (**Fig. 6D-E**). Smokers exhibited a higher proportion of clonal mutations but a lower fraction of binding mutations, suggesting a greater degree of immunoediting in clonal neoantigens. This provides a potential explanation for the weaker performance of the clonality-aware neoantigen landscape in homogeneous NSCLC tumors, as the remaining clonal neoantigens may have survived due to low immunogenicity or immune evasion mechanisms, thereby weakening their predictive value.

## DISCUSSION

Computational approaches to prioritize neoantigens remain an important area of development to provide biomarkers of immunotherapy response and provide insights into the characteristics of mutated peptides that govern immunogenicity^69,70^. Our immunogenicity model, NeoPrecis-Immuno, enhances predictive performance by incorporating MHC-binding motifs, refining how peptides are represented in relation to TCR recognition. The allele benefit score derived from this model further demonstrates significant predictive power for patient outcome in ICI treatment. In addition, we integrated both MHC-I and MHC-II predictions to improve ICI response prediction, recognizing the cooperative roles of CD8+ and CD4+ T cells in anti-tumor immunity. By further incorporating tumor clonality, we improved response prediction in melanoma and provided a more nuanced description of heterogeneous tumors in NSCLC.

Peptide binding to MHC molecules is a critical determinant of pMHC-TCR interactions, traditionally thought to be driven primarily by non-anchor residues that directly contact the TCR. However, emerging evidence suggests that MHC molecules influence TCR recognition beyond their role in peptide presentation. Wu *et al.* demonstrated that MHC polymorphisms can induce conformational changes in pMHC complexes, affecting TCR-pMHC interactions and modulating immune activation^71^. Additionally, modifications to MHC-binding anchor residues have been shown to alter TCR recognition. Cole *et al.* found that anchor-modified peptides could activate T cells with distinct TCRs^72^, while Smith *et al.* showed that anchor residues function as allosteric modulators, shaping pMHC-TCR interactions^73^. These findings align with our model interpretation, where MHC-specific motif enrichment refines the embedding space at different scales, particularly capturing the high variability of anchor residues, ultimately improving neoantigen immunogenicity prediction.

Beyond amino acid substitutions and MHC molecules, which influence TCR recognition, we hypothesize that the entire peptide sequence also contributes to immunogenicity. Richman *et al.* demonstrated that dissimilarity to the human proteome correlates with peptide immunogenicity^74^, while Łuksza *et al.* introduced a “non-selfness” metric based on neoantigen similarity to known antigens^31^. In our model, we attempted to capture peptide-level immunogenicity using a sigmoid-scaled cross-reactivity distance, allowing the model to learn how peptide sequence influences immune recognition. However, component contribution analysis showed that the sigmoid scaling factor had minimal impact on prediction performance (**Fig. 3A**), likely due to limited sample size of the training set. The human proteome consists of millions of peptides, yet our model was trained on only thousands, potentially restricting its ability to distinguish self from non-self effectively. More targeted modeling of peptide selfness may be necessary to further enhance immunogenicity predictions.

In the context of ICI response evaluation, while MHC-I-restricted neoantigens have traditionally been the primary focus in mutation burden analysis, growing evidence underscores the essential role of MHC-II in tumor immunity. Magen *et al.* demonstrated that ICI response correlates with the clonal expansion of both intratumoral CD4+ and CD8+ T cells in hepatocellular carcinoma patients^75^. Similarly, Espinosa-Carrasco *et al.* emphasized the necessity of the immune triad— CD4+ T cells, CD8+ T cells, and dendritic cells—for effective tumor elimination^43^. These findings align with our results, showing that combining MHC-I and MHC-II predictions for a given mutation improves ICI response prediction.

Tumor heterogeneity remains a major obstacle to effective ICI responses^34–38^, underscoring the need to identify patients with heterogeneous tumors who may derive therapeutic benefit. The NeoPrecis clonality-aware neoantigen landscape, NeoPrecis-LandscapeClone, significantly improved response prediction in heterogeneous NSCLC tumors. However, its performance was notably weaker in homogeneous NSCLC tumors, where dominant clones with a lower proportion of binding mutations likely underwent immunoediting reducing their impact on immunogenicity (**Fig. 6D-E**). This is supported by Rosenthal *et al.*^76^, who demonstrated that clonal mutations in early-stage NSCLC may undergo immunoediting, suggesting their role in shaping response to ICI. Nevertheless, this phenomenon is not observed in melanoma, which has a higher ratio of clonal mutations and a greater proportion of binding mutations (**Fig. S9A-B**). Niknafs *et al.*^77^ indicated key differences in clonal persistent TMB between melanoma and NSCLC, showing that melanoma exhibits a stronger correlation between immune response and clonal persistent TMB, aligned with our observations. These findings suggests that tumor type-specific adaptations are necessary to optimize neoantigen landscape modeling.

While NeoPrecis significantly enhances the characterization of neoantigen immunogenicity, it has certain limitations. First, the model is currently not applicable to indels or frameshift mutations, as it requires a defined wild-type peptide sequence. However, these mutations can generate highly immunogenic neoantigens and should be considered in future extensions^78^. Additionally, the model focuses solely on the core binding region, whereas MHC-II has been shown to involve flanking regions that contribute to both binding affinity and immunogenicity^79,80^. Furthermore, key immunogenic determinants, including germline variations, the tumor microenvironment, and T-cell infiltration, are not explicitly modeled. Sayaman *et al.*^81^ and Pagadala *et al.*^82^ demonstrated that germline predisposition variants can shape the tumor microenvironment, affecting immune infiltration and, consequently, tumor immunity. Future efforts should integrate these immune and tumor-intrinsic factors to further refine neoantigen prediction and improve personalized immunotherapy strategies.

## METHODS

### Preparation of the TCR binding triplet dataset

TCR-peptide binding data were obtained from IEDB^48^ and VDJdb^49^. Binding complexes were included if they contained a complete annotation of the beta complementarity-determining region 3 (CDR3), peptide sequence, and MHC allele. To isolate TCR recognition from the effects of MHC binding, peptides with low MHC-binding affinity were removed using a %rank threshold of 2 for MHC-I and 10 for MHC-II, based on predictions from NetMHCpan-4.1^51^ (MHC-I) and NetMHCIIpan-4.3^52^ (MHC-II). After filtering, the dataset included 69,278 complexes for MHC-I and 3,165 complexes for MHC-II. Peptide-MHC complexes were then grouped by MHC allele and CDR3 sequence, ensuring that all peptides within a group bound to the same MHC and TCR molecule. Groups containing only one peptide were removed.

To train the immunogenicity model to distinguish mutated peptides from wild-type peptides, we constructed a dataset of peptide triplets. Each triplet is consisted of a seed peptide (a reference peptide), a positive peptide (similar to the seed), and a negative peptide (dissimilar to the seed) (**Fig. S1A**). The model was trained to minimize the distance between seed-positive pairs (similar peptides) while maximizing the distance between seed-negative pairs (dissimilar peptides). To mimic the effect of single amino acid substitutions, the difference between both seed-positive and seed-negative pairs was constrained to one Hamming distance. Seed-positive pairs were sampled from each peptide-MHC complex group to ensure functional similarity between the seed and positive peptides. All possible pairs were sampled, yielding 1,153 MHC-I and 261 MHC-II seed-positive pairs.

For each seed-positive pair, a negative peptide was randomly selected to be one Hamming distance away from the seed but absent from the positive set, which comprised peptides within the same peptide-MHC complex group. A BLOSUM62 matrix was used as a prior, ensuring that the substitution between seed and negative resulted in a negative BLOSUM62 score, indicating substantial sequence dissimilarity. To enhance dataset diversity, we applied 10-fold generation, ultimately producing 11,530 MHC-I and 2,610 MHC-II triplets (**Table S1**).

### Preparation of the CEDAR immunogenicity dataset

Cancer-related immunogenicity data were obtained from CEDAR. Samples were included if they contained annotations for the MHC allele, wild-type peptide, mutated peptide, and immunogenicity label (T-cell assay). To ensure accurate modeling of substitutions, samples where the wild-type and mutated peptides had mismatched lengths were removed. To isolate TCR recognition from the effects of MHC binding, peptides with low binding affinity were filtered out using a %rank threshold of 2 for MHC-I and 10 for MHC-II. After filtering, the dataset comprised 4,176 MHC-I samples and 87 MHC-II samples (**Table S2**). Then, the dataset was split into training (75%), validation (10%), and testing (15%) sets. The testing set initially contained 640 samples, from which 438 unique peptides—ensured to be unobserved by any of the predictors—were retained for evaluation. Due to the limited number of MHC-II samples, this restriction was not applied to MHC-II.

### Preparation of the NCI cohort

The NCI gastrointestinal cancer cohort, derived from Parkhurst *et al.*^53^, was utilized to evaluate the immunogenicity model and construct a multi-dimensional integrated model. The cohort comprises 7,384 missense mutations across 75 patients, with both MHC-I and MHC-II alleles available for all patients, along with abundance data (DNA allelic fraction, RNA allelic fraction, and RNA expression level). Notably, RNA expression levels were quartile-normalized for each individual. Each mutated peptide underwent T-cell assay testing, resulting in mutation-centric data with labels for each mutation. This approach differs from peptide-centric data (e.g., CEDAR dataset) where each peptide has a label. In total, 41 mutations exhibited CD8+ activity, and 34 showed CD4+ activity. For peptide-centric features, such as MHC-binding affinity and immunogenicity scores, an aggregation approach was applied to obtain mutation-centric predictions. This method will be described in detail in a subsequent section.

To evaluate immunogenicity prediction, mutations were filtered based on abundance and MHC presentation metrics to exclude factors that could bias TCR recognition comparison. Mutations with zero DNA allelic fraction, zero RNA allelic fraction, or no expression level were removed. Additionally, mutations were filtered using a %rank threshold of 2 for MHC-I and 10 for MHC-II. The final dataset for immunogenicity prediction evaluation consisted of 1,088 mutations for MHC-I (36 positives) and 1,336 mutations for MHC-II (30 positives). For the construction and evaluation of the multi-dimensional integrated model, all 7,384 mutations were included to account for all three dimensions of the analysis (**Table S3**).

### NeoPrecis immunogenicity model

The NeoPrecis immunogenicity model, NeoPrecis-Immuno, is a custom machine learning framework that integrates key immunogenicity determinants, including residue-level embedding, MHC-binding motif enrichment, peptide embedding, geometric representation, and sequence-aware scaling (**Fig. 2a**). pMHC complexes are first predicted using an MHC-binding predictor, and the 9-mer core binding region is identified as the input sequence. NeoPrecis-Immuno encodes amino acids using the BLOSUM62 matrix, followed by projection into a two-dimensional latent space via a single-layer neural network, capturing amino acid relationships in the context of TCR recognition. The model then refines embeddings by incorporating MHC-binding motifs through position-wise projection, accounting for the complex interactions between pMHC and TCR. Peptide embeddings are aggregated using position-weighted pooling, emphasizing structurally important core residues. To differentiate wild-type and mutated peptides, NeoPrecis-Immuno employs a geometric representation, highlighting sequence alterations relevant to immune recognition. A two-layer neural network then computes a scaling factor for geometric distance, generating the cross-reactivity distance (CRD), ensuring that sequence composition differences influence immunogenicity prediction. Finally, CRD and MHC-binding predictions are integrated into a single-layer neural network, producing the final neoantigen immunogenicity score.

The immunogenicity model was trained in two stages. In the first stage, it was trained on the binding triplet dataset using contrastive learning with triplet loss to optimize CRD:

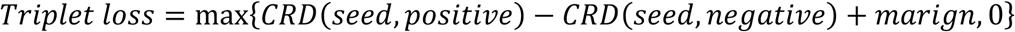

where *CRD(m, n)* represents the cross-reactivity distance between peptides *m* and *n*, and the *margin* was set to 1. In the second stage, the model was fine-tuned on the CEDAR immunogenicity dataset using binary cross-entropy loss to predict neoantigen immunogenicity scores. During fine-tuning, all layers were frozen, except for the scaling factor and classifier, allowing the model to focus on adjusting immunogenicity predictions. Training was conducted for 100 epochs with a learning rate of 0.005, using the AdamW optimizer.

### Residue embeddings

We utilized three distinct residue embeddings: BLOSUM62 embedding, NeoPrecis-Immuno residue embedding, and motif-enriched embedding. To generate BLOSUM62 embeddings, we applied principal component analysis (PCA) to the BLOSUM62 matrix, extracting the top two principal components as the embedding for each amino acid. NeoPrecis-Immuno residue embedding is also a two-dimensional representation designed to capture amino acid relationships in the context of TCR recognition. The motif-enriched embedding is a position-wise transformation of the NeoPrecis-Immuno residue embedding, incorporating position-specific MHC-binding motifs. Each position-specific motif is represented as a projection matrix, which is then multiplied with the residue embedding to refine its representation based on MHC-binding context. This transformation ensures that amino acid embeddings reflect not only intrinsic residue properties but also their contextual influence within the pMHC complex.

### Immunogenicity predictions from other predictors

To benchmark NeoPrecis-Immuno, we evaluated its performance against three immunogenicity predictors: PRIME^30^, ICERFIRE^27^, and DeepNeo^28^. All three models provide MHC-I predictions, but only DeepNeo supports MHC-II. However, due to differences in MHC allele coverage, not all predictors were available for every allele in our datasets. To ensure fair comparisons in the CEDAR internal validation, we restricted the testing set to peptides supported by all predictors. For the NCI external validation, if a predictor did not support certain alleles in a patient, we removed those alleles and computed the mutation-centric immunogenicity score based on the remaining available alleles. Predictions were aggregated across alleles using the masked maximum approach. If a mutation lacked predictions from a given predictor across all its binding alleles, it was excluded from the testing set to maintain consistency in benchmarking.

For PRIME, we used PRIME 2.0, downloaded from GitHub. ICERFIRE was run as a stand-alone application, excluding expression values from its input. DeepNeo predictions were obtained using the DeepNeo-v2 web server, which only supports the beta chain of MHC-II (DQB, DPB). Consequently, alpha chains (DQA, DPA) were ignored when computing mutation-centric DeepNeo scores. These standardized procedures ensured a fair and unbiased evaluation of NeoPrecis-Immuno against existing immunogenicity predictors.

### Component contribution analysis

To assess the contribution of individual components in the NeoPrecis-Immuno, we systematically extracted and analyzed key features, including residue embedding, motif enrichment, position weighting, and CRD scaling factors. The model first computes substitution distance (SubDist), which uses residue embeddings to measure differences between wild-type and mutated peptides. This is then refined into position-weighted substitution distance (SubPosDist), which adjusts SubDist by incorporating positional importance, ensuring that residues within the peptide-binding core are weighted according to their functional contribution to TCR recognition. Further refinement is achieved through geometric distance (GeoDist), which extends SubPosDist by integrating motif enrichment. By incorporating MHC-binding motifs, GeoDist adjusts residue embeddings to better reflect TCR-pMHC interactions. Finally, cross-reactivity distance (CRD) applies a sigmoid scaling function to GeoDist, modifying geometric distances to account for sequence-specific variations in immunogenicity. This scaling ensures that even when geometric distances between wild-type and mutated peptides are similar, differences in sequence composition are considered in immunogenicity predictions.

### Affine transformation for the motif-enriched embedding

To interpret how motif enrichment refines residue embeddings, we applied an affine transformation between the original residue embedding and the motif-enriched embedding:

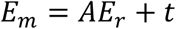

where *E_m_* is the motif-enriched embedding, *E_r_* is the original residue embedding, *A* is the transformation matrix, and *t* is the translation vector. To analyze how this transformation scales the embedding space, we performed singular value decomposition (SVD) on *A*:

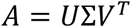

where *Σ* contains the scaling factors, and *V^T^* provides the principal directions of transformation. The scaling effect along the original x- and y-axes was computed by projecting *Σ* onto the original coordinate system using *V^T^*:

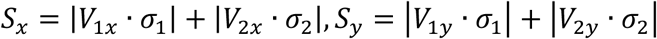

where *S_x_* and *S_y_* represent the scaling along the x- and y-axes, and *v_1_*, *v_2_* are the basis vectors from *V^T^*. This analysis quantifies how motif enrichment reshapes residue embeddings, refining their representation in the context of MHC-binding motifs.

### Allele benefit scores

Allele benefit scores are designed to evaluate how MHC-binding motifs refine neoantigen immunogenicity by influencing T-cell recognition. For each allele and position, we calculated scaling factors (*S_x_* and *S_y_*), and the allele benefit score was defined as the average geometric mean of *S_x_* and *S_y_* across all 9-mer positions:

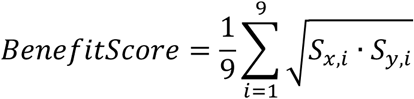

A higher allele benefit score results from larger *S_x_* or *S_y_* values, indicating a greater distance between WT and MT peptides, which in turn suggests increased immunogenicity. Additionally, the allele benefit score can be aggregated across individual alleles to provide an overall assessment of an individual’s immune response. The geometric mean was calculated separately for MHC-I and MHC-II alleles, where *N* is the number of alleles:

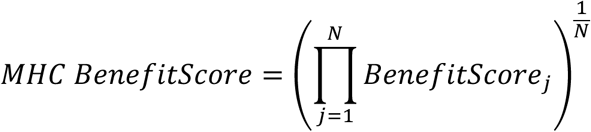

The MHC-dual benefit score was computed as the geometric mean of the MHC-I and MHC-II benefit scores:

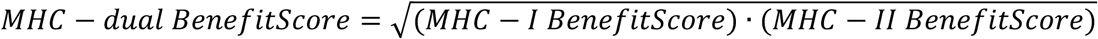

To integrate with TMB, the combined score was calculated as the product of *logTMB* and MHC-dual benefit score:

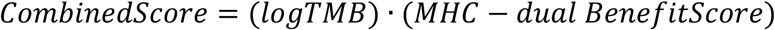

### Calculation of multi-dimensional metrics

Eight metrics were computed to evaluate neoantigen immunogenicity, categorized into abundance, presentation, and recognition metrics. Abundance metrics included DNA allele frequency (AF), RNA AF, and RNA expression quartile, while presentation metrics comprised robustness and PHBR. Recognition metrics included agretopicity ratio, foreignness, and NeoPrecis-Immuno. Abundance metrics were available in the NCI cohort, whereas for the Ott and ICI cohorts, DNA AF was derived from variant calling results, and RNA AF was calculated using bam-readcount^83^. RNA expression quartile was computed as the quartile transformation of TPM values obtained from RSEM^84^ for each individual. Presentation metrics were determined as follows: robustness was defined as the number of binding alleles, using a %rank threshold of 2 for MHC-I and 10 for MHC-II. PHBR (patient harmonic-mean best rank) was computed as the harmonic mean of the %rank of the best-binding peptide for each allele. Recognition metrics included agretopicity ratio, which was calculated as the ratio of MHC binding scores between mutated and wild-type peptides, and foreignness, which measured sequence similarity to foreign immunogenic antigens. Foreignness was computed using the NeoantigenEditing repository (GitHub) by Łuksza *et al.*^31^

### T-cell recognition metric aggregation across MHC alleles

To assess the impact of MHC allele aggregation on immunogenicity prediction, we tested various approaches. The first consideration was masking, which excludes alleles that do not bind to the neoantigen based on the %rank threshold of 2 for MHC-I and 10 for MHC-II. The second consideration was the choice of aggregation method, where we evaluated mean, harmonic mean, weighted average (weighted by binding affinity), best binding, and maximum. For example, the masked maximum method selects the highest score among all binding alleles that meet the masking criteria. This approach ensures that only strongly binding alleles contribute to the aggregated immunogenicity score, refining the prediction of neoantigen immunogenicity.

### Feature selection for the integrated multi-dimensional model

To identify the most predictive features among the eight immunogenicity metrics, we applied recursive feature elimination with cross-validation (RFECV) using five-fold cross-validation. Features were eliminated one at a time, and the subset yielding the best cross-validation performance was selected as the final feature set. For consistency, we retained five features for both MHC-I and MHC-II, even though the optimal feature set for MHC-II consisted of only two features. This decision was made because there was no significant difference in performance between using two and five features in MHC-II (**Fig. S5A**), and the smaller sample size for MHC-II limited the reliability of feature selection.

### Integrated multi-dimensional model

The integrated model, NeoPrecis-Integrated, is a logistic regression trained on the selected features, with all features undergoing standard normalization based on the training set. The mean and standard deviation calculated from the training set were recorded and subsequently applied to the testing set for consistent normalization. To assess the impact of multi-dimensional integration, we conducted five-fold cross-validation on the NCI dataset. For ICI response prediction, the integrated model was trained on the entire NCI dataset to ensure robust performance.

### Preparation of the ICI cohorts

We compiled eight patient cohorts treated with immune checkpoint inhibitors (ICIs), including five melanoma cohorts and three NSCLC cohorts (**Table S6**). In total, 695 patients (443 melanoma and 252 NSCLC) with whole-exome sequencing (WES) data were collected from dbGaP. Raw sequencing data were processed using a standardized pipeline. Variant calling was performed using Nextflow-Sarek (v3.4.2)^85,86^, with the hg38 reference genome, employing Mutect2 for somatic variant calling, VEP for mutation annotation, and ASCAT for copy number analysis, tumor purity, and ploidy estimation. RNA sequencing data were aligned to hg38 using STAR (v2.7.3a)^87^ with default settings, and transcript quantification was conducted using RSEM (v1.3.1)^84^. HLA typing was performed with HLA-HD (v1.7.0)^88^, ensuring accurate identification of patient-specific MHC alleles. Clonality analysis was conducted using PyClone (v0.13.1)^65^, with 5,000 iterations and a maximum of 100 clusters, allowing for a detailed assessment of tumor heterogeneity.

### Calculation of mutation burden

We computed three distinct mutation burden metrics for comparison: tumor mutation burden (TMB), tumor neoantigen burden (TNB), and NeoPrecis-Immuno burden (NPB). TMB was defined as the total number of non-synonymous mutations in the tumor. TNB accounted for non-synonymous mutations that met an MHC-I binding threshold (PHBR ≤ 2), incorporating MHC-binding constraints into the burden calculation. NPB incorporated both MHC-I and MHC-II binding along with TCR recognition. A mutation was counted if the product of NeoPrecis-Immuno-I (MHC-I) and NeoPrecis-Immuno-II (MHC-II) scores was ≥ 0.16. This threshold was derived from the median NeoPrecis-Immuno score (around 0.4) for both MHC-I and MHC-II, where 0.16 represents the squared value (0.4 × 0.4), ensuring that mutations with strong immunogenic potential were captured.

### Calculation of tumor-centric immunogenicity score

Aggregating neoantigen immunogenicity scores into a tumor-centric immunogenicity score is essential for assessing individual immune response. To achieve this, we first computed mutation-centric immunogenicity scores using three distinct metrics: NeoPrecis-Immuno-I, representing the NeoPrecis-Immuno score for MHC-I; NeoPrecis-Immuno-II, representing the NeoPrecis-Immuno score for MHC-II; and NeoPrecis-Immuno-dual, calculated as the product of NeoPrecis-Immuno-I and NeoPrecis-Immuno-II, thereby integrating both MHC-I and MHC-II pathways. Unless otherwise specified, NeoPrecis-Immuno-dual was used as the default metric in our experiments of ICI response prediction.

We explored three methods to aggregate these mutation-centric scores into a comprehensive tumor-centric immunogenicity score. The NeoPrecis-LandscapeSum method involved summing the NeoPrecis-Immuno scores across all identified neoantigens, providing a straightforward cumulative measure. The NeoPrecis-LandscapeCCF approach leveraged the cancer cell fraction (CCF) by applying a weighted sum based on mutation prevalence as inferred from PyClone analysis, thus incorporating the tumor’s cellular composition into the immunogenicity assessment. The NeoPrecis-LandscapeClone method integrated tumor clonality to offer a more nuanced perspective. Using PyClone, mutations were grouped into clusters according to their prevalence in the tumor and mutation size. Within each cluster, NeoPrecis-Immuno scores were summed to derive cluster-specific immunogenicity, which was then averaged across clusters, weighted by their prevalence, to calculate the tumor-centric immunogenicity score (**Fig. 6A**). The three NeoPrecis-Landscape scores were log-transformed to facilitate interpretation within a more manageable range of values.

### Clonality-aware approaches for benchmarking

We benchmarked NeoPrecis-Landscape against two other approaches that derive information from tumor clonality, the Cauchy-Schwarz index of neoantigens (CSiN)^66^ and the immunoediting-based optimized tumor neoantigen load (ioTNL)^67^.

The CSiN score is calculated by aggregating the normalized product of variant allele frequency (VAF) and neoantigen load across various quality cutoffs for predicted peptide-MHC (pMHC) binding. For each mutation, we considered all possible 8 to 11-mer (MHC-I) and 15-mer (MHC-II) mutant peptides paired with their respective MHC alleles. The neoantigen load (*L_i_*) for a given mutation *i* represents the number of these pMHC pairs that surpass a specific MHC-binding prediction rank percentile cutoff (*c*). Following the original CSiN methodology, we employed rank percentile cutoffs of 0.375, 0.5, 0.625, 0.75, 1.25, 1.75, and 2. Mutations with a VAF (*V_i_*) below 0.05 were excluded. For samples with over 500 mutations, only the 500 mutations with the highest VAF were retained for analysis. The CSiN score is then computed using the following equation:

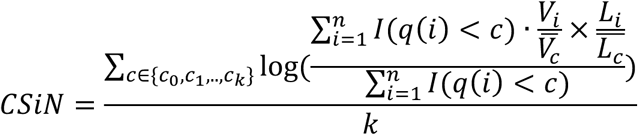

where *k* is the total number of cutoffs used, *n* is the number of mutations considered for the patient, and *q(i)* is the best rank percentile among all pMHCs for mutation *i*.

The ioTNL score is calculated by summing the product of cancer cell fraction (*CCF*) and neoantigen load for subclones identified through PyClone analysis^65^ that exhibit evidence of immune elimination. An immune elimination score (*L_j_/S_j_*) is calculated for each subclone to quantify its potential for immune elimination; only subclones with a score below a defined cutoff (*e*) are included. The ioTNL score is computed using the following equation:

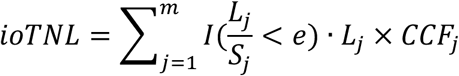

where *m* is the total number of subclones, *L_j_* is the neoantigen load in subclone *j*, *S_j_* is the number of nonsynonymous mutations in subclone *j*, and *e* is the predefined cutoff for the immune elimination score. The selection of the cutoff (*e* = 1.4) was empirically determined to maximize the predictive performance of ioTNL across our datasets. Although this data-driven approach carries a risk of overfitting and potential information leakage, no established biological rationale was available to guide this selection.

### Survival analysis

For survival analysis, patients were stratified into two groups based on their target metric scores, with one group comprising the top half and the other comprising the bottom half of the distribution. Due to differences in data availability, we used overall survival (OS) for melanoma and progression-free survival (PFS) for NSCLC. Kaplan-Meier survival curves were generated, and log-rank tests were performed to assess statistical differences between the two groups.

### Tumor heterogeneity assessment

To evaluate tumor heterogeneity, we calculated the Gini index at two levels: cluster mutation size and cluster prevalence, capturing diversity in both dimensions. The Gini index quantifies inequality, with a value of 0 indicating perfect equality (all clusters have equal mutation counts or prevalence) and a value of 1 indicating maximum inequality (a single cluster contains all mutations or has a prevalence of 1). The Gini index (*G*) is computed as follows, where *n* represents the number of clusters and *x* denotes either mutation size or prevalence:

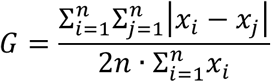

To classify tumors into homogeneous and heterogeneous groups, we applied a two-dimensional approach based on prevalence Gini index (pGini) and size Gini index (sGini). For each cancer type, tumors with pGini ≤ median and sGini ≤ median were categorized as heterogeneous, while tumors with pGini > median and sGini > median were classified as homogeneous.

## Acknowledgements

ICI cohorts were collected from several published studies. The melanoma cohort from Hugo and colleagues was obtained from Sequence Read Archive (SRA) under accessions SRP090294 and SRP067938. The melanoma cohorts from Van Allen *et al.* and Liu *et al.* were obtained from dbGaP under accession phs000452, supported by the National Human Genome Research Institute (NHGRI) Large Scale Sequencing Program, Grant U54 HG003067 to the Broad Institute (PI, Lander). The melanoma cohort from Riaz and colleagues was obtained from SRA under accession SRP094781. The melanoma cohort from Snyder and colleagues was obtained from dbGaP under accession phs001041; we thank Martin Miller at Memorial Sloan Kettering Cancer Center (MSKCC) for his assistance with the NetMHC server, Agnes Viale and Kety Huberman at the MSKCC Genomics Core, Annamalai Selvakumar and Alice Yeh at the MSKCC HLA typing laboratory for their technical assistance, and John Khoury for assistance in chart review. The NSCLC cohort from Rizvi and colleagues was obtained from dbGaP under accession phs000980. We thank the members of the Thoracic Oncology Service and the Chan and Wolchok labs at MSKCC for helpful discussions. We thank the Immune Monitoring Core at MSKCC, including L. Caro, R. Ramsawak, and Z. Mu, for exceptional support with processing and banking peripheral blood lymphocytes. We thank P. Worrell and E. Brzostowski for help in identifying tumor specimens for analysis. We thank A. Viale for superb technical assistance. We thank D. Philips, M. van Buuren, and M. Toebes for help performing the combinatorial coding screens. The data presented in this paper are tabulated in the main paper and in the supplementary materials. This work was supported by the Geoffrey Beene Cancer Research Center (MDH, NAR, TAC, JDW, AS), the Society for Memorial Sloan Kettering Cancer Center (MDH), Lung Cancer Research Foundation (WL), Frederick Adler Chair Fund (TAC), The One Ball Matt Memorial Golf Tournament (EBG), Queen Wilhelmina Cancer Research Award (TNS), The STARR Foundation (TAC, JDW), the Ludwig Trust (JDW), and a Stand Up To Cancer-Cancer Research Institute Cancer Immunology Translational Cancer Research Grant (JDW, TNS, TAC). Stand Up To Cancer is a program of the Entertainment Industry Foundation administered by the American Association for Cancer Research. The NSCLC cohort from Anagnostou and colleagues was obtained from dbGaP under accession phs001940, supported in part by US National Institutes of Health grant CA121113. The NSCLC cohort from Ravi and colleagues was obtained from dbGaP under accession phs002822. We express our deep gratitude to the patients and families whose participation enabled this study. We further thank the respective sequencing centers at Yale University, Johns Hopkins University, and the Broad Institute of MIT and Harvard for processing the whole exome and RNA-seq data presented here. Funding for this study was provided by a Stand Up To Cancer - American Cancer Society Lung Cancer Dream Team Translational Research Grant (Grant Number: SU2C-AACR-DT17-15). Stand Up to Cancer is a program of the Entertainment Industry Foundation. Research grants are administered by the American Association for Cancer Research, the scientific partner of SU2C. This work was additionally supported by The Mark Foundation for Cancer Research (Grant Number: 19-029-MIA) Expanding Therapeutic Options for Lung Cancer (EXTOL) project.

## Author Contributions

Conceptualization: KHL, TJS, MZ, HC. Methodology: KHL. Investigation: KHL, TJS. Visualization: KHL. Funding acquisition: HC. Project administration: HC. Supervision: MZ, HC. Writing – original draft: KHL. Writing – review & editing: TJS, MZ, HC.

## Data and Materials Availability

Peptide-centric TCR-pMHC binding data are downloaded from the Immune Epitope Database (IEDB, https://www.iedb.org) and VDJdb (https://vdjdb.cdr3.net). Immunogenicity data are downloaded from the Cancer Epitope Database and Analysis Resource (CEDAR, https://cedar.iedb.org). Mutation-centric immunogenicity data (NCI dataset) are obtained from Parkhurst *et al.* (https://doi.org/10.1158/2159-8290.CD-18-1494). ICI cohorts are obtained with the following accession numbers: Hugo *et al.* (SRP090294, SRP067938), Van Allen *et al.* (SRP011540), Snyder *et al. (*SRP072934), Riaz *et al.* (SRP094781), Liu *et al.* (SRP011540), Ravi *et al.* (SRP413932), Anagnostou *et al.* (SRP238904), Rizvi *et al.* (SRP064805). Codes for conducting immunogenicity prediction and neoantigen landscape evaluation are deposited at https://github.com/cartercompbio/NeoPrecis.

## Supplementary Figures

**Fig. S1.**
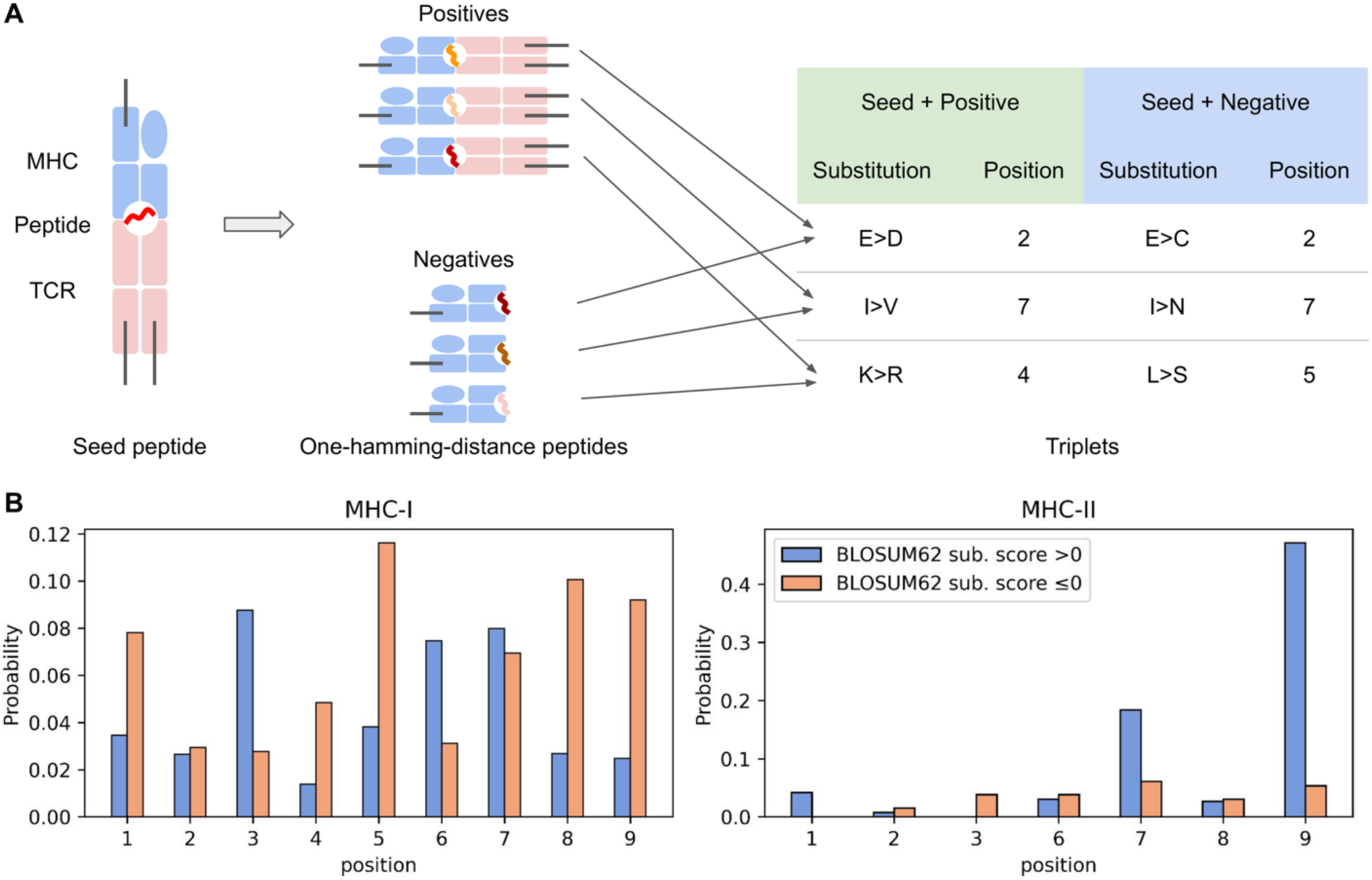
Preparation and characteristics of the TCR-binding triplet dataset. a. A schematic diagram of the triplet generation. For each TCR-peptide complex serving as the seed, positive samples were identified by selecting remaining peptides that share the same MHC allele and CDR3 sequence but differ by one Hamming distance in the peptide sequence. For each seed-positive pair, a negative sample was randomly selected as a peptide with a one-Hamming-distance difference from the seed that was not included in the positives. Each triplet consists of a seed, a positive, and a negative sample. b. Frequency of amino acid substitutions grouped by BLOSUM62 substitution scores and positions in seed-positive pairs (mimicking non-immunogenic substitutions) for MHC-I and MHC-II.

**Fig. S2.**
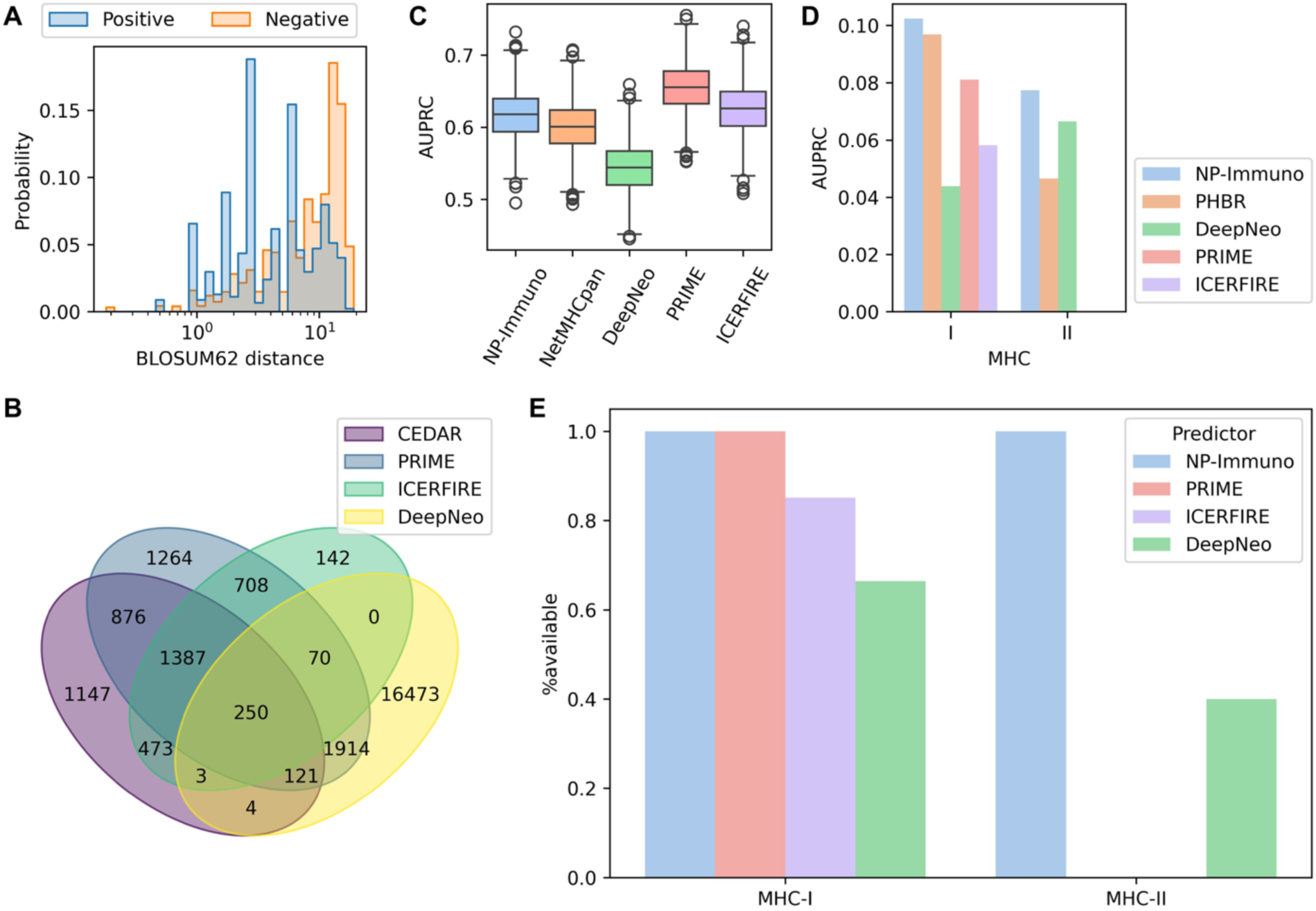
Validation of NeoPrecis (NP)-Immuno. **(A)** Distribution of BLOSUM62 distances in the TCR-binding triplet dataset. The positive group consists of positive pairs (seed and positive peptides) that mimic non-immunogenic substitutions, while the negative group consists of negative pairs (seed and negative peptides) that mimic immunogenic substitutions. **(B)** Venn diagram showing the overlap between the CEDAR dataset, PRIME training set, ICEFIRE training set, and DeepNeo training set for MHC-I. **(C)** Area under the precision-recall curve (AUPRC) for each predictor on MHC-I samples (n=438) from the internal testing set (CEDAR dataset). AUROC distributions were calculated using bootstrapping (k=1,000). In the boxplots, the center line represents the median, boxes indicate the interquartile range (IQR; 25th–75th percentile), and whiskers extend to the most extreme data points within 1.5× the IQR. Outliers are depicted as individual circles. NP denotes NeoPrecis. **(D)** AUPRC comparisons of each predictor on MHC-I (n=1,448) and MHC-II (n=1,373) samples from the external testing set (NCI dataset). **(E)** Allele availability across predictors in the NCI cohort. The y-axis represents the proportion of alleles covered by each predictor.

**Fig. S3.**
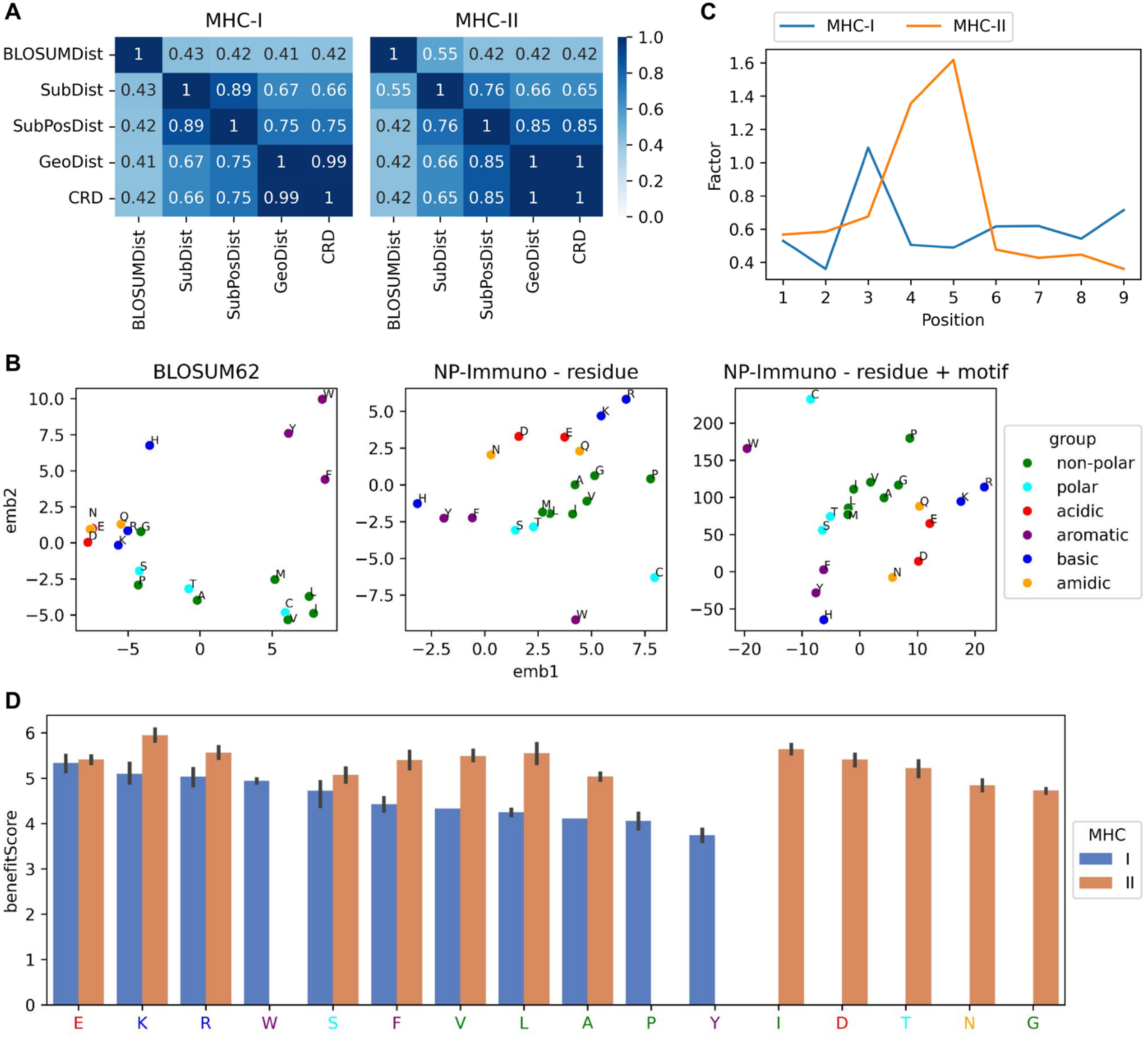
Interpretation of NeoPrecis (NP)-Immuno. **(A)** Correlation between components for MHC-I and MHC-II. **(B)** Residue embedding transformation from BLOSUM62, model embedding, to motif-enriched embedding, illustrated using the second position of allele B*40:01 as an example. **(C)** Position factors for MHC-I and MHC-II. **(D)** Distribution of allele benefit scores, with alleles grouped by the dominant amino acid in the anchor motif.

**Fig. S4.**
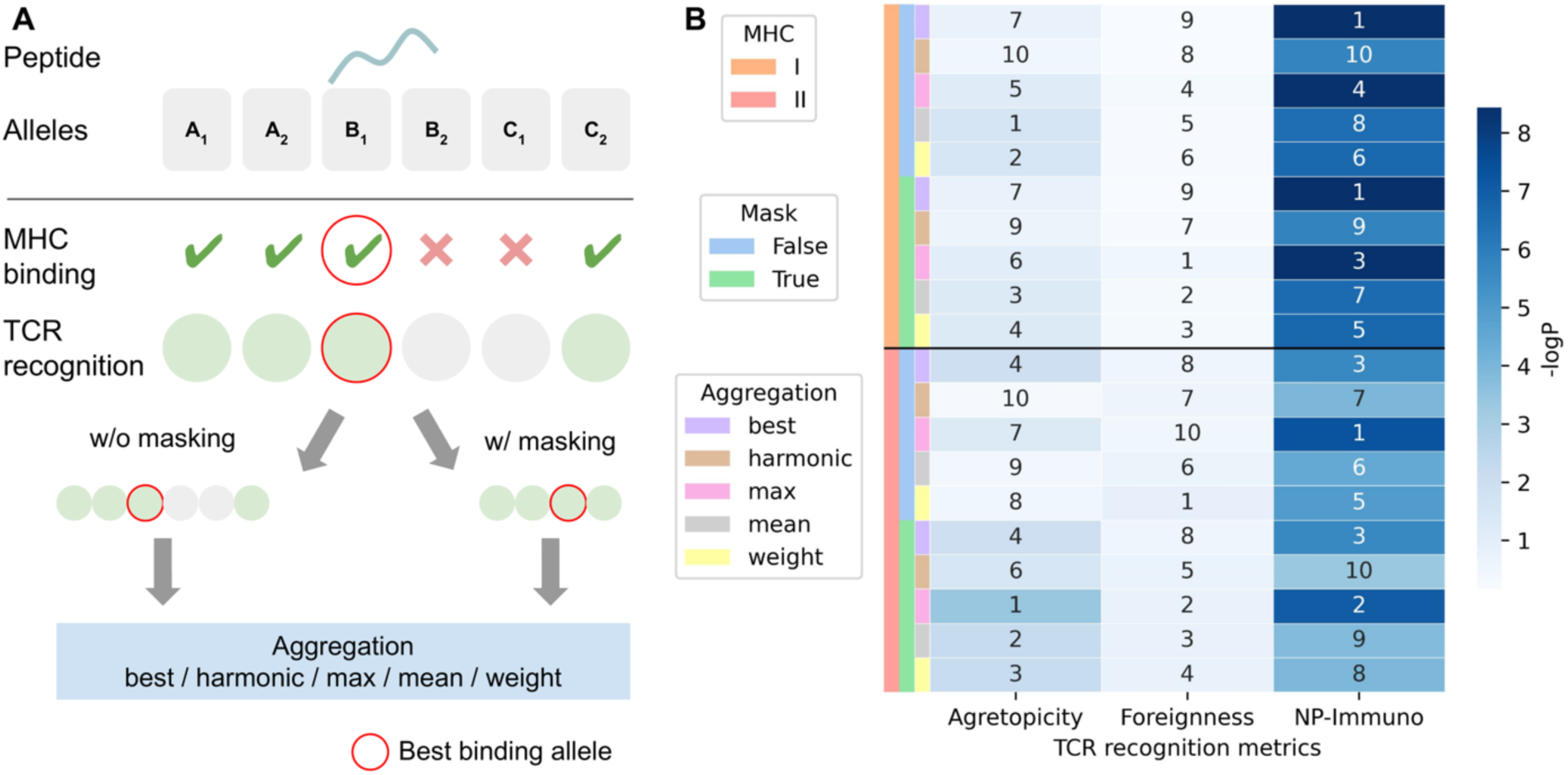
Comparison of various aggregation approaches. **(A)** The schematic illustrates the allele aggregation process. First, MHC-binding and TCR recognition metrics are calculated for each allele, and the best-binding allele is identified. In the masking approach, alleles without MHC binding are excluded, whereas in the non-masking approach, all alleles are retained. Various aggregation methods are then applied: “Best” represents the recognition score of the best-binding allele, “harmonic” represents the harmonic mean of recognition scores, “max” represents the highest value, “mean” represents the average, and “weight” represents the weighted average, with weights based on binding affinity. **(B)** The heatmap displays the negative log p-values from the association tests (Mann–Whitney U test) between each approach and T-cell activity across various TCR recognition metrics. Annotations indicate the rank within each MHC class and for each metric.

**Fig. S5.**
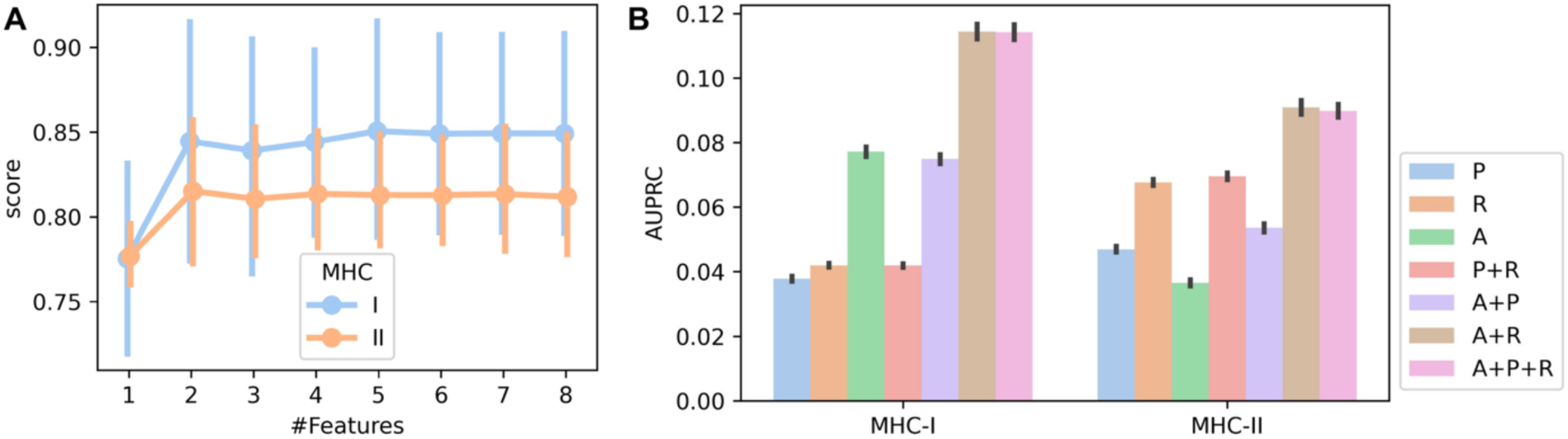
Feature selection and performance evaluation of NeoPrecis-Integrated. **(A)** Results of recursive feature elimination with cross-validation (RFECV) for selecting features for MHC-I and MHC-II. **(B)** Area under the precision-recall curve (AUPRC) for cross-validation on the NCI cohort using different combinations of metrics.

**Fig. S6.**
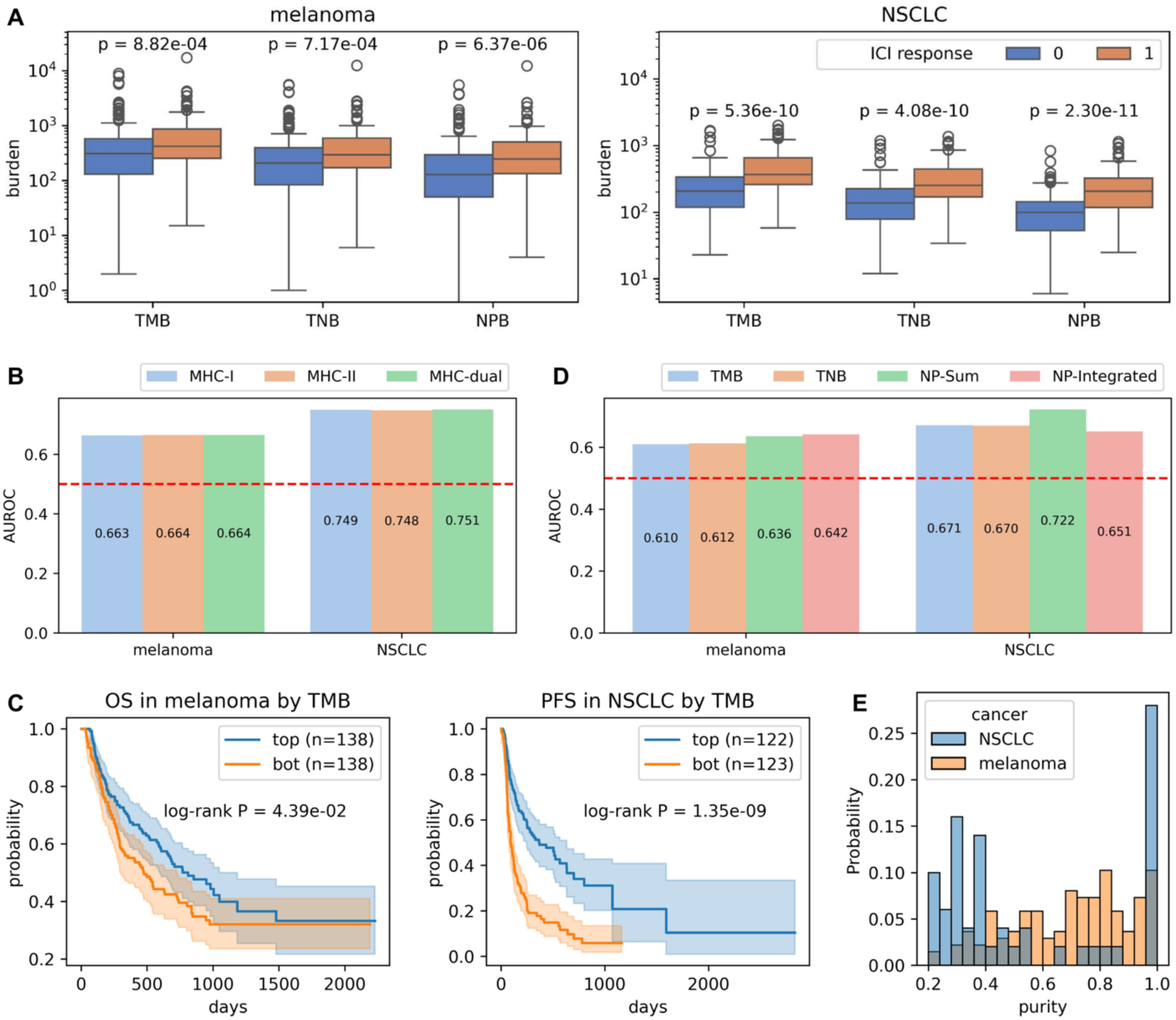
Performance evaluation of NeoPrecis on ICI response prediction. **(A)** Comparison of mutation burden distributions for melanoma and NSCLC. P-values were calculated using the Mann-Whitney U test. **(B)** AUROC comparison of NeoPrecis-LandscapeSum scores calculated using MHC-I-only, MHC-II-only, or combined MHC-I and MHC-II (dual). **(C)** Kaplan-Meier survival curves stratified by TMB, with log-rank P-values shown for melanoma and NSCLC. **(D)** AUROC comparison of TMB, TNB, NeoPrecis-LandscapeSum (NP-Sum), and NeoPrecis-Integrated (NP-Integrated) on RNA-available samples (n = 187; 137 melanoma and 50 NSCLC). **(E)** Purity distribution of RNA-available samples in melanoma and NSCLC.

**Fig. S7.**
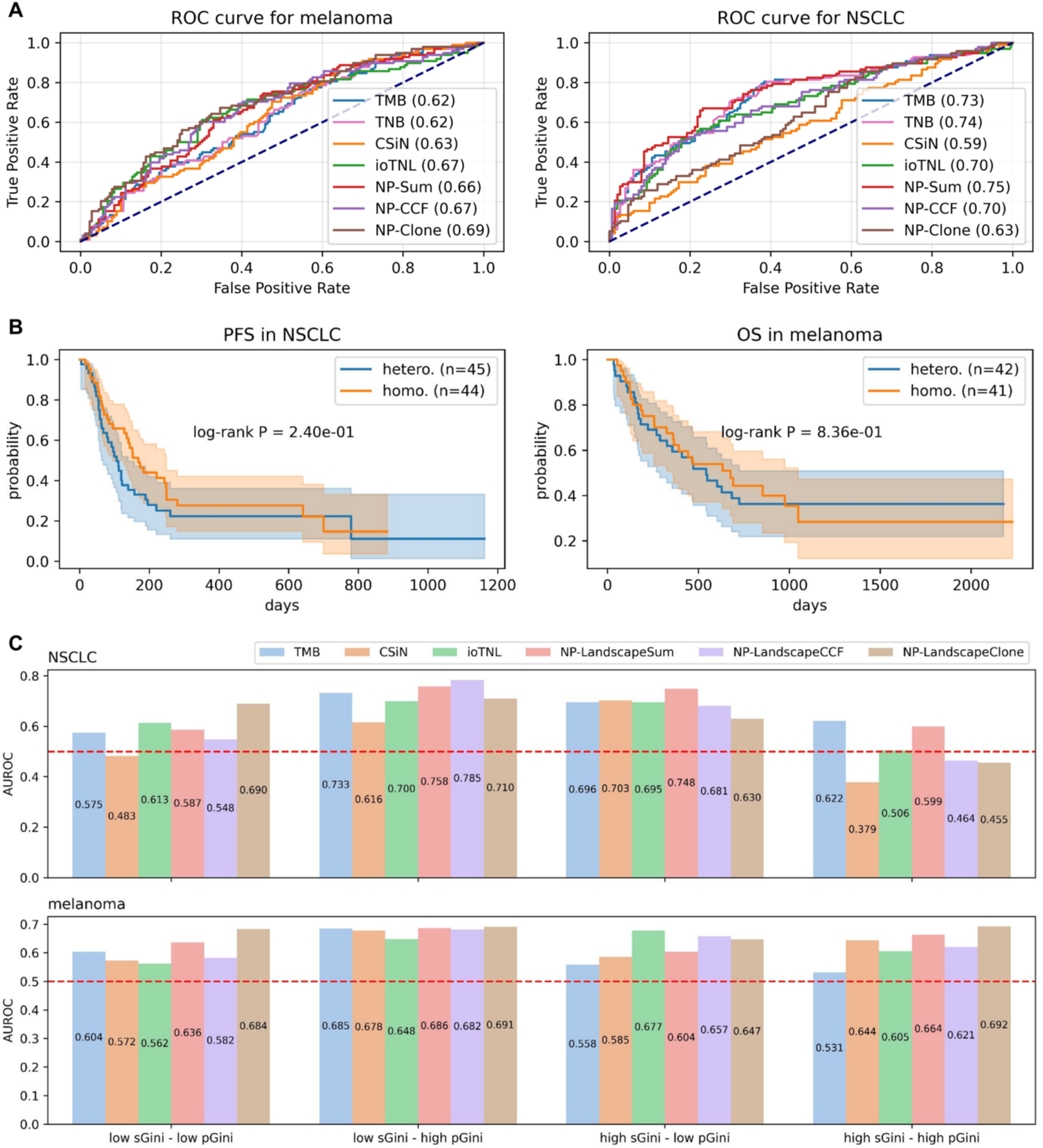
Clonality analysis in melanoma and NSCLC. **(A)** AUROC comparison of tumor-centric immunogenicity metrics, including TMB, TNB, CSiN, ioTNL, NeoPrecis-LandscapeSum (NP-Sum), NeoPrecis-LandscapeCCF (NP-CCF), and NeoPrecis LandscapeClone (NP-Clone), for response prediction in melanoma and NSCLC. **(B)** Survival curves comparing homogeneous (homo.; high sGini–high pGini) and heterogeneous (hetero.; low sGini–low pGini) tumor groups in melanoma (OS) and NSCLC (PFS). **(C)** AUROC comparison of TMB, CSiN, ioTNL, NP-LandscapeSum, NP-LandscapeCCF, NP-LandscapeClone across four patient subgroups in melanoma and NSCLC. NP denotes NeoPrecis.

**Fig. S8.**
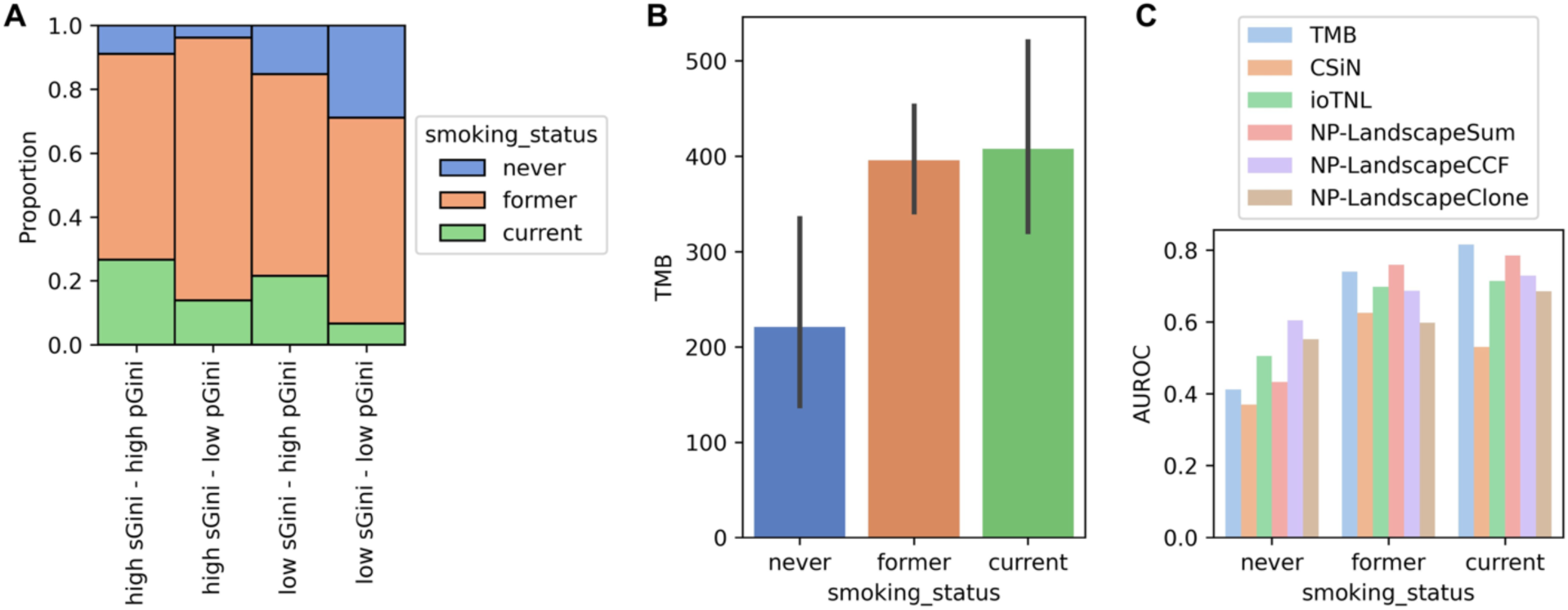
Smoking status in NSCLC. **(A)** Distribution of smoking status across four heterogeneity-defined subgroups. **(B)** TMB distribution by smoking status. **(C)** AUROC comparison of TMB, CSiN, ioTNL, NP-LandscapeSum, NP-LandscapeCCF, and NP-LandscapeClone across different smoking statuses. NP denotes NeoPrecis.

**Fig. S9.**
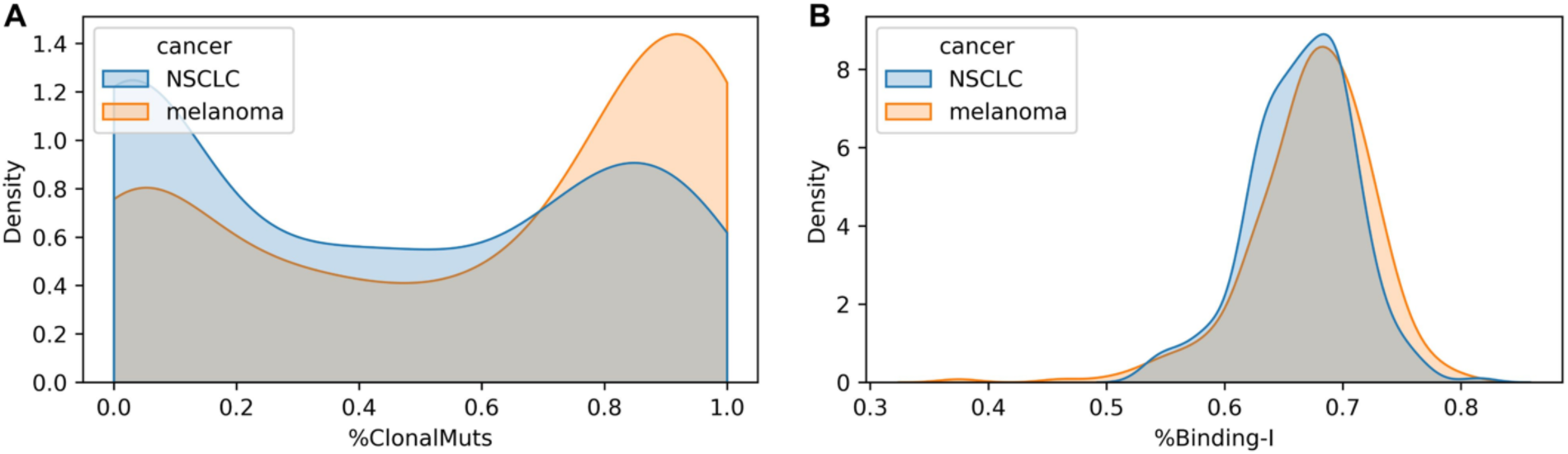
Immunoediting analysis across cancer types. **(A)** Density plot estimated using KDE showing the distribution of %ClonalMuts, the ratio of clonal mutations (CCF ≥ 0.85) to total mutations, across melanoma and NSCLC. Values are clipped between 0 and 1. **(B)** Density plot estimated using KDE showing the distribution of %Binding-I, the ratio of MHC-I binding mutations (PHBR ≤ 2) to total mutations, across melanoma and NSCLC. Values are clipped between 0 and 1.

## Supplementary Tables

Table S4 and S6 are presented below, while Table S1-3, S5, and S7 are in separated .csv files.

**Table S1. Dataset of TCR binding triplets.**

Each row in this dataset represents a TCR binding triplet, comprising a seed peptide, a positive peptide, and a negative peptide. The seed and positive peptides are single Hamming distance variants that bind to the same MHC molecule and TCR CDR3 region, as identified from the TCR binding data obtained from IEDB and VDJdb. Negative peptides, each one Hamming distance from the seed but not present in the TCR binding data, were randomly selected. To ensure MHC binding, these negative peptides were subjected to binding prediction. For each seed-positive pair, five negative peptides with the same mutated position and five with different mutated positions were sampled. In total, we produced 11,530 MHC-I and 2,610 MHC-II triplets.

**Table S2. CEDAR immunogenicity dataset.**

Each row in this dataset represents a cancer-related mutant peptide with an immunogenicity label verified by T-cell assays, derived from the CEDAR dataset. For both wild-type and mutated peptides, MHC binding predictions were calculated. Additionally, immunogenicity predictions from PRIME, DeepNeo, ICERFIRE, and NeoPrecis-Immuno were included, along with an annotation indicating their presence in each predictor’s training set.

**Table S3. NCI gastrointestinal cancer cohort dataset.**

This dataset, derived from Parkhurst et al., includes 7,384 missense mutations across 75 patients. Abundance annotations (DNA allelic fraction (DNA_AF), RNA allelic fraction (RNA_AF), and RNA expression quartile level (RNA_EXP_QRT)) and immunogenicity labels (CD4 and CD8 T-cell activations) were obtained from the original study. For each mutation, we calculated MHC presentation metrics (robustness and PHBR) and T-cell recognition metrics (agretopicity ratio, foreignness, and NeoPrecis-Immuno). Additionally, benchmark scores from PRIME, DeepNeo, and ICERFIRE were incorporated.

**Table S5. Allele benefit scores.**

The benefit score, derived from the scaling factors of NeoPrecis-Immuno for each allele, is a mutation-independent metric representing the potential for enhanced tumor immunity associated with that allele.

**Table S7. Immune checkpoint inhibitor cohort dataset.**

Each row in this dataset represents a patient, with metadata including age, sex, ICI response, and survival data. Mutation burden, CSiN, ioTNL, NeoPrecis prediction, and clonality analysis results are also annotated for each patient.

**Table S4.** Components of the NeoPrecis immunogenicity model. To evaluate the contribution of individual components to NeoPrecis-Immuno, we assessed model performance by systematically including or excluding key features: residue embedding, position factors, motif enrichment, and sigmoid scaling. BLOSUM62 embedding served as a baseline for comparison. The rows of the table represent the different model configurations evaluated, while the columns indicate the presence or absence of each key component in a given model.

|  | BLOSUM62<br>embedding | Residue<br>embedding | Position<br>factors | Motif<br>enrichment | Sigmoid<br>scaling |
| --- | --- | --- | --- | --- | --- |
| BLOSUMDist | v |  |  |  |  |
| SubDist |  | v |  |  |  |
| SubPosDist |  | v | v |  |  |
| GeoDist |  | v | v | v |  |
| CRD |  | v | v | v | v |

**Table S6.** Meta data of the ICI cohorts. Each row in this table represents a cancer cohort from a specific study, annotated with the cancer type (Cancer), the total number of patients (#Samples), the number of patients with RNA-seq data available (#RNA), and the number of patients who responded to immune checkpoint inhibitors (#Response).

|  | Cancer | #Samples | #RNA | #Response |
| --- | --- | --- | --- | --- |
| Hugo et al. | melanoma | 38 | 27 | 21 |
| Van Allen *et al.* | melanoma | 110 | 40 | 17 |
| Riaz et al. | melanoma | 109 | 84 | 24 |
| Snyder *et al.* | melanoma | 64 | 0 | 36 |
| Liu et al. | melanoma | 122 | 122 | 50 |
| Rizvi *et al.* | NSCLC | 34 | 0 | 12 |
| Anagnostou *et al.* | NSCLC | 36 | 0 | 17 |
| Ravi et al. | NSCLC | 182 | 51 | 68 |

